# Elucidating design principles for Ribozyme-Enabled Tissue Specificity (RETS) to allow precise expression without specialized promoters

**DOI:** 10.1101/2025.08.14.670357

**Authors:** Max M. Combest, Josh Conlin, Vivia Van De Mark, Morgan Boisen, Hadley Colwell, Arjun Khakhar

## Abstract

Tissue- or cell type- specific expression of transgenes is often essential for interrogation of biological phenomenon or predictable engineering of multicellular organisms but can be stymied by cryptic enhancers that make identification of promoters that generate desired expression profiles challenging. In plants the months-to-years long timeline associated with prototyping putative tissue-specific promoters in transgenic lines deepens this challenge. We have developed a novel strategy called *Ribozyme Enabled Tissue Specificity* (RETS) that leverages the knowledge of where and when genes are expressed derived from transcriptomic studies to enable tissue-specific expression without needing characterized promoters. It uses a split self-splicing ribozyme based on a group I intron from *Tetrahymena thermophila* to enable conditional reconstitution of a transgene mRNA in the presence of a secondary tissue-specific mRNA of choice. We elucidate the design features that enable flexible swapping of transgenes and targets, enhancing transgene expression, and circumventing host RNA interference responses. We then show that these innovations enable tissue-specific and dose-dependent expression of transgenes in *Arabidopsis thaliana*. Finally, we demonstrate the utility of RETS both for creating genetically encoded biosensors to study the spatiotemporal patterns of gene expression *in planta* as well as for engineering tissue-specific changes in organ size. RETS provides a novel avenue to study expression patterns of native loci with non-destructive imaging, complementing the weakness of existing approaches. Additionally, the spatiotemporal control of transgene expression afforded by RETS enables precision engineering of plant phenotypes which will facilitate enhancing crops without the trade-offs associated with constitutive expression.

## Introduction

Precision spatiotemporal control of transgene expression is often critical for both interrogating gene function as well as engineering novel traits in plants (Bull & Khakhar, 2023; Hong et al., 2003; Maizel & Weigel, 2004). The expression of transgenes in inappropriate cell and tissue types can cause embryonic lethality or generate negative pleiotropic phenotypes that confound experiments or engineering goals (Bandopadhyay et al., 2010). Modern transcriptomics has generated temporally resolved maps of native gene expression at tissue and single-cell resolution across a range of plant species (He et al., 2023; Lim et al., 2022). These data have been leveraged to identify genes with desired expression profiles and their promoters have been used to enable tissue-specific transgene expression (Tian et al., 2023). However, identifying the sequence necessary to recapitulate a transcripts expression pattern is often stymied by the presence of cryptic enhancers that can be embedded in introns or present hundreds of kilobases away from the gene (Hummel et al., 2023; Tan et al., 2023; Tian et al., 2022; Ullah et al., 2018; Yasmeen et al., 2023). Additionally, the limited existing understanding of regulatory elements (Tian et al., 2022, 2023; Yasmeen et al., 2023) makes it challenging to reliably predict if these promoters will generate consistent expression patterns across different genomic insertion points, genetic backgrounds, and environmental conditions. This often necessitates iterative prototyping in stable transgenic lines to identify novel tissue-specific promoters, which severely limits the utility of this approach thanks to the months- to years-long timeline associated with plant transgenesis (Bandopadhyay et al., 2010). Thus, there is a pressing need for tools that can leverage transcriptomic data to enable spatiotemporal control of transgene expression without the need to find and characterize new promoters. Recent advances in engineering catalytic RNA motifs, called ribozymes, provide a promising avenue to achieve this goal.

Since the discovery of self-splicing group I introns, groups have been working on ways to engineer them to facilitate targeted *trans-*splicing (Gambill et al., 2023; Hasegawa et al., 2004, 2006; Hasegawa & Rao, 2006; Kruger et al., 1982; Sullenger & Cech, 1994; Watanabe & Sullenger, 2000; Zaug et al., 1986). A group I intron from the rRNA of *Tetrahymena thermophila* functions as a ribozyme (TtRz), catalyzing its own excision and the ligation of its bordering exons without the need for any protein partners (Zaug & Cech, 1986). Recently, Gambill et al. developed an RNA detection platform based on these motifs in bacteria (Gambill et al., 2023) wherein it was embedded in the coding sequence of a reporter. This transcript was then split into two at the first loop of the ribozyme, producing two separate RNA molecules, which were extended with guide sequences that are complementary to a chosen target RNA. When the guides bind to their target, they scaffold the two portions of the TtRz close enough to reform a functional structure and allow for the splicing reaction to occur, ligating the neighboring exons and thereby reconstituting the reporter coding sequence to enable expression (Figure 1). This mechanism represents an ideal starting point to enable tissue-specific expression without tissue-specific promoters.

**Figure 1.**
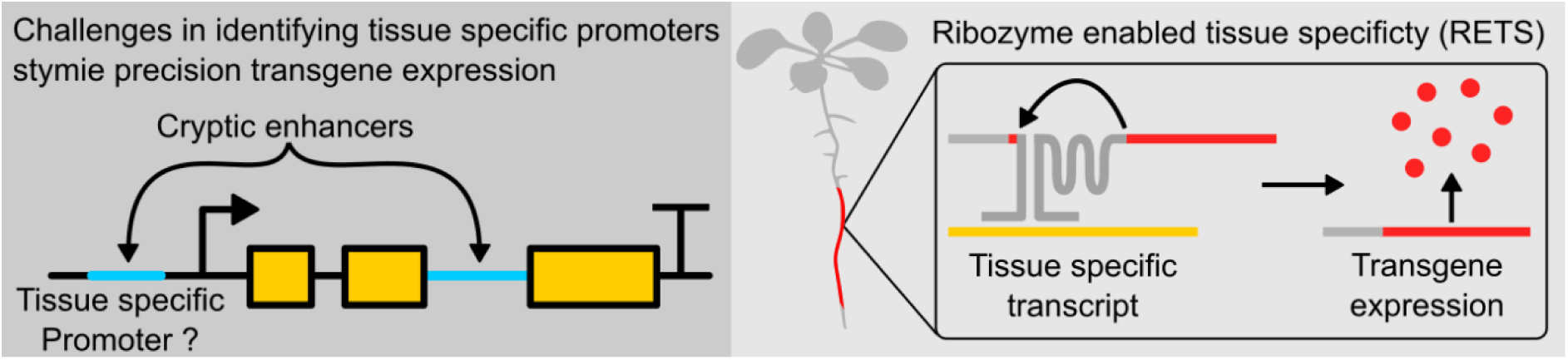
Graphical abstract. (Left) Tissue-specific expression can be complicated by cryptic enhancer sequences that make identifying tissue-specific promoters challenging. (Right) Ribozyme Enabled Tissue Specificity, RETS, allows tissue-specific expression of transgenes without the need for characterizing tissue-specific promoters.

In this work we have developed a promoter-independent tissue-specific expression system in plants called *Ribozyme Enabled Tissue Specificity* (RETS). Here, a chosen cargo is divided into two inactive transcripts that bear split ribozyme sequences fused to guide sequences that are complementary to a tissue-specific transcript of choice. The two transcripts are expressed constitutively, but the protein cargo is directed to be expressed only in those tissues where the target RNA is available to scaffold the ribozyme and promote splicing-based reconstitution of the translatable mRNA, thereby producing tissue-specific expression without the need for tissue-specific promoters. We use transient expression assays in *Nicotiana benthamiana* to elucidate design features that enable flexible swapping of transgenes and targets, enhance transgene expression, and circumvent host RNA interference (RNAi) responses. We go on to demonstrate that the system can drive tissue-specific and dose-dependent expression of transgenes in stable transgenic lines of the model plant *Arabidopsis thaliana*. Finally, we demonstrate the utility of RETS both for creating genetically encoded biosensors to study the spatiotemporal patterns of gene expression *in planta* as well as for engineering tissue-specific changes in organ size.

## Results

### The RETS system enables modular RNA-gated gene expression *in planta*

We first aimed to test whether the RNA-templated reconstitution of a split self-splicing ribozyme that was previously demonstrated in bacteria (Gambill et al., 2023) could be modified to be functional *in planta*. We constructed a transfer DNA (T-DNA) plasmid encoding constitutive expression cassettes for the two halves of a split fluorescent reporter with the split ribozymes targeted to a test transcript, the coding sequence of an *Arabidopsis*-codon-optimized Cas9, as well as a second fluorescent protein, Venus, to enable ratiometric normalization to control for differential delivery. Here the split reporter, a red fluorescent protein (mScarlet), was split at the U-base of the start codon, AUG (Figure 2A). We reasoned that this modular split site within the start codon would enable the reporter to be easily swapped with other proteins in the future. To enable transcript dependent reconstitution, the two halves of the ribozymes were fused to 200-base (bp) long guide arms that had sequence complementarity with two adjacent regions on the Cas9 transcript. A detailed description of the construct design can be found in the methods section and supplementary materials (Supplemental RETS construct design). If this system functioned correctly, we would expect mScarlet signal only in the presence of the Cas9 transcript. From here forward, RETS constructs will be described with the notation, *Target*:RETS:Cargo, so this first construct that expresses mScarlet in response to Cas9 is referred to as “*Cas9*:RETS:mScarlet”.

**Figure 2.**
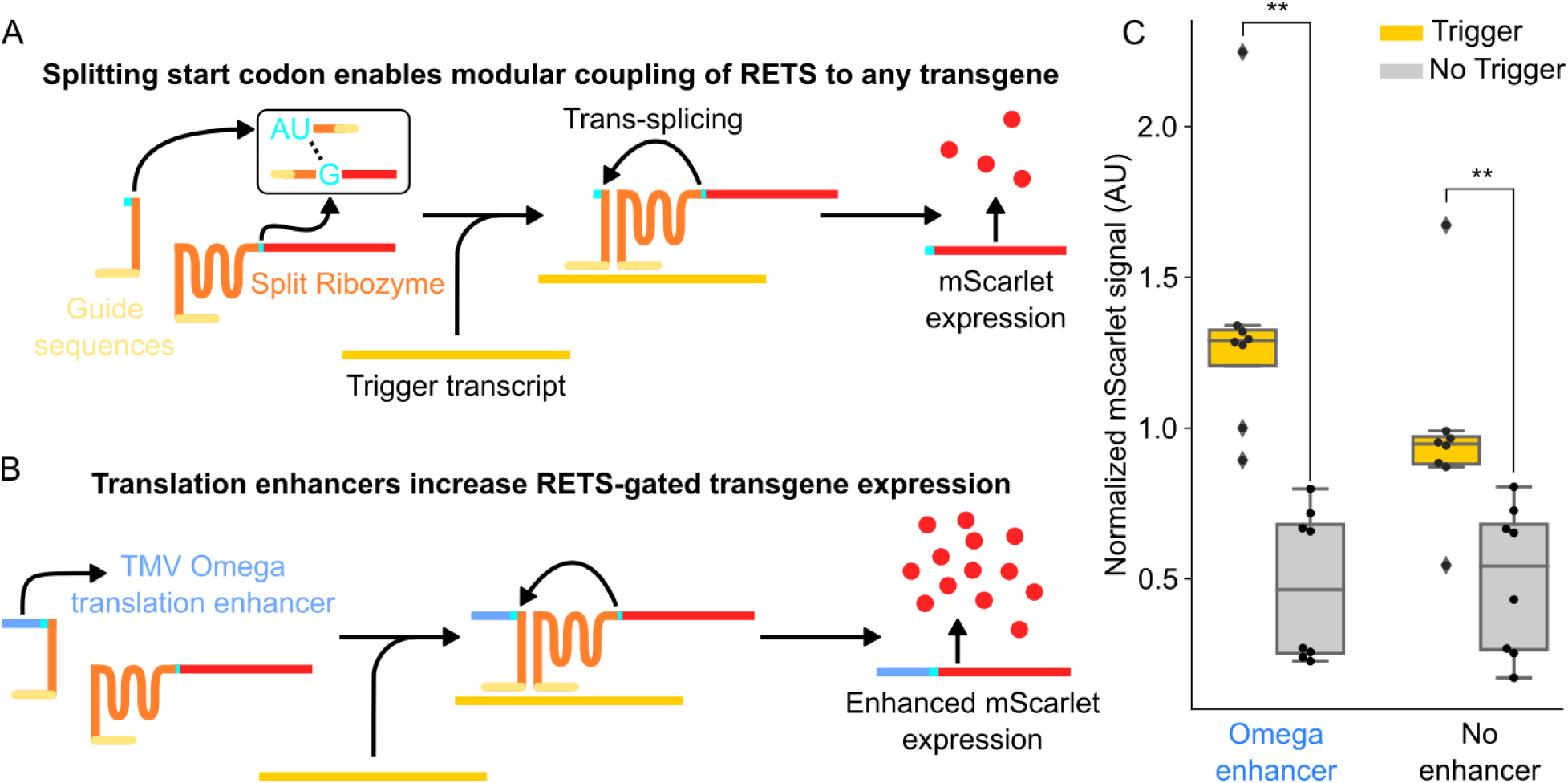
RETS conditions cargo expression on presence of target RNA. A) Schematic describing the RETS mechanism. Here the type I intron ribozyme (orange) is split with each section fused to either the AU or G of the start codon of an mScarlet reported (red). Additionally, the other side of both sections are fused to guide sequences (light yellow) with sequence complementarity with a trigger mRNA (dark yellow). Annealing to this trigger brings the two halves of RETS together enabling splicing dependent reconstitution and expression of mScarlet. B) Schematic showing how the TMV Omega enhancer (blue) can be incorporated 5’ of the split start codon to enhance mScarlet output. C) Boxplots summarizing mScarlet signal normalized by Venus signal from transient fluorescent assays of the constructs show in A (No enhancer) and B (Omega enhancer) with (gray) or without (yellow) co-infiltration with the trigger transcript. Each dot represents an independent biological replicate and ** represents p ≤ 0.005.

To test *Cas9*:RETS:mScarlet, we used *Agrobacterium* infiltration (Kapila et al., 1997) to transiently express this construct, either with or without co-infiltration of a second T-DNA expressing the Cas9 transcript, in the leaves of *Nicotiana benthamiana*. At 96hrs post infiltration, we observed that the ratiometrically normalized mScarlet output was significantly higher when the Cas9 transcript was present than when it was absent, with an average fold change of 1.97x (p=0.004) (Figure 2C). We further hypothesized that the fold change of RETs could be enhanced via the inclusion of a translational enhancer to generate more output signal from each correctly spliced transcript (Figure 2B). To test this, we included a strong translational enhancer, the omega leader sequence from tobacco mosaic virus(Gallie, 2002), upstream of the start codon and tested it in tandem with the original construct using the assay described above. At 96hrs post infiltration, the ratiometrically normalized mScarlet fold change of the system with the omega sequence in the presence of Cas9, at 2.78x (p=0.0003) (Figure2C), was significantly greater than the one lacking it. Based on these results, all future constructs incorporated this enhancer. Taken together, these results demonstrate that RETS is functional *in planta* and that its output can be enhanced via the incorporation of translational enhancers.

### RETS enables tissue-specific expression of transgenes without needing tissue-specific promoters

Building on the transient validation of the RETS mechanism, we set out to test the utility of the RETS system to enable tissue-specific transgene expression, by mimicking the expression pattern of native genes, in stable transgenic lines. As a proof-of-concept we chose to use the *Arabidopsis* sucrose transporter gene, *SUCROSE-PROTON SYMPORTER 1* (*SUC1),* which has been demonstrated by prior studies to be expressed in trichomes (Lasin et al., 2020; Sivitz et al., 2006, 2008). These studies showed that *SUC1*’s expression pattern is dependent on the presence of enhancer elements in its introns and so cannot be duplicated by reporters using only its promoter (Lasin et al., 2020). This makes it a good demonstration of the promoter-independent functionality of the RETS system. We constructed a new T-DNA expressing a modified version of the previously described RETS construct, with guide sequences that target the first exon of At*SUC1*, and used them to generate transgenic *SUC1:*RETS:mScarlet *A. thaliana* lines. The reporter expression across several independent lines was subsequently characterized using fluorescence microscopy. We observed that mScarlet fluorescence was localized to the bases and cell bodies of young trichomes of rosette leaves (Figure 3B-D, See Figure S1A to compare to untransformed Col-0) as expected (Lasin et al., 2020; Sivitz et al., 2006). These results demonstrate that the RETS system can program transgene expression to mimic the expression pattern of native genes without needing tissue-specific promoters. It also shows how the RETS system can be flexibly reprogramed to target a transcript of choice by swapping guide sequences.

**Figure 3.**
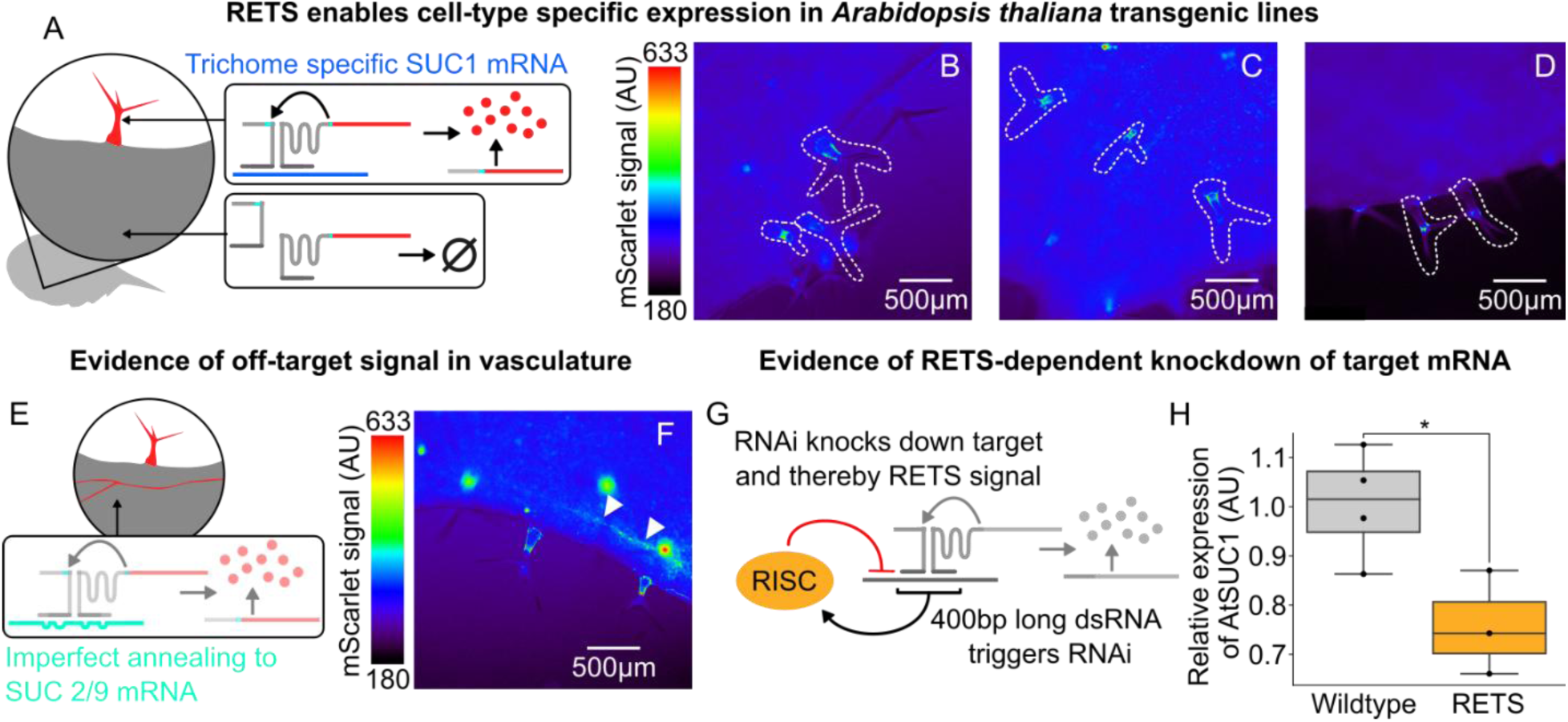
RETS enables tissue-specific expression of cargo with some caveats for 200nt guide sequences. A) Cartoon describing how RETS would restrict cargo expression to trichomes due to trichome specific expression of SUC1 (blue). B-D) Representative fluorescence microscopy images of mScarlet signal from adult leaves of SUC1:RETS:mScarlet expressing plants with the trichomes highlighted by dashed white lines. E) Cartoon describing how unexpected mScarlet fluorescence in vascular tissues could be driven by cross-reactivity to vascularly localized members of the sucrose transporter family, SUC2 and SUC9 (teal). F) Representative fluorescence microscopy images of mScarlet signal from adult leaves of SUC1:RETS:mScarlet expressing plants with mScarlet signal in vasculature highlighted with white arrows. G) Cartoon depicting how the production of dsRNA intermediates during the process of target recognition could trigger RNA silencing directed at the target and/or the RETS cassettes. H) Box plots summarizing RT-PCR expression data for AtSUC1 relative to a housekeeping gene in wildtype (gray) and SUC1:RETS:mScarlet expressing plants (orange). Each dot represents a biological replicate and * corresponds to p ≤ 0.05.

Unexpectedly, while characterizing these lines we also observed expression of mScarlet throughout the vasculature of adult leaves where *SUC1* is not expressed (Figure 3F). We hypothesized that this might be due to non-specific binding of the guide sequences to non-target transcripts leading to reconstitution of mScarlet. An alignment of the 400 bases of *SUC1* targeted by the guide sequences with other members of the SUC transporter family revealed a significant (∼75%) sequence similarity to vascularly localized(Sivitz et al., 2006) members, namely *SUC2* and *SUC9* (Figure S2A). This incomplete homology is may be sufficient to trigger the RETS mechanism and highlights the need for transcript specific guide sequences. Additionally, we hypothesized that the transient formation of duplex RNA during the association of the RETS transcripts with the target could act as a trigger for RNAi, potentially downregulating the target and both RETS transcripts, and leading to a reduction in mScarlet signal as well as unwanted pleiotropies (Figure 3G). To assess whether this was occurring we measured the expression of the target, *SUC1*, as well as a housekeeping gene to enable normalization in both *SUC1*:RETS:mScarlet lines and wildtype *A. thaliana* seedlings via RT-qPCR. We observed that the RETS lines had consistently lower levels of *SUC1* expression (mean decrease of 32.5%, p=0.034) (Figure 3H) than wildtype seedlings indicating a silencing effect. Taken together, these results show that a lack of guide sequence specificity and silencing both limit the functionality of RETS for enabling tissue-specific expression.

### Reducing guide length improves fold-change of RETS and reduces silencing

Seeking to resolve the design issues identified by characterization of the *SUC1*:RETS:mScarlet, we chose to modify the length and target of the guide sequences (Figure 4A). To reduce the chances of inducing an RNAi response from RNA duplex formation we redesigned new RETS constructs that have shorter guides of either 30, 25 or 20 bp each. These lengths were chosen as prior work has shown that RNA duplexes under 37 bp in length reduce the efficiency of DICER based generation of siRNA (Kakiyama et al., 2019). The functionality of these shorter guide sequences was tested via transient expression experiments in *N. benthamiana*, either with or without co-infiltration with a constitutively expressed copy of *AtSUC1* as the trigger, using the ratiometrically normalized reporter assay described above. We observed that all three guide lengths generated significantly higher normalized mScarlet signal in the presence of the trigger compared to when it was absent, demonstrating all three guide lengths were functional. Additionally, we observed an increased fold change in mScarlet output for the shortened guides compared to the previously test long (200bp) guides, ranging from 2.3-4.5x for each of the shortened guide sequences compared to 1.8 for 200 base guides (Figure S3). Interestingly we observed that one of the constructs with the shortest guide length (20bp), generated the largest fold change in reporter signal (4.5x, p = 3.4×10^−8^, Figure S3). We next made versions of the shortened-guide construct that targeted a more unique section of the At*SUC1* transcript to reduce off-target activity. We used the NCBI BLAST search tool (Sayers et al., 2024) to identify a region that has no significant similarity to any other endogenous transcript. These were also tested using the same transient expression assay in *N. benthamina* described above. Here again we observed that they retained the capacity to reconstitute the mScarlet transcript in the presence of the *SUC1* transcript and that they had a higher fold change in response to the target than the original RETS construct with 200 bp long guides. Overall, we observed that the RETS constructs with shorter guides outperformed the 200bp version both in terms of absolute activation by the target and fold change. For the shortened guides targeting the unique sequence, fold increase in mScarlet output for each construct varied between 1.7 and 2.0 (Figure S3). Among these, the best performing was the 30-base guide construct targeting the unique region of SUC1 (Figure 4B).

**Figure 4.**
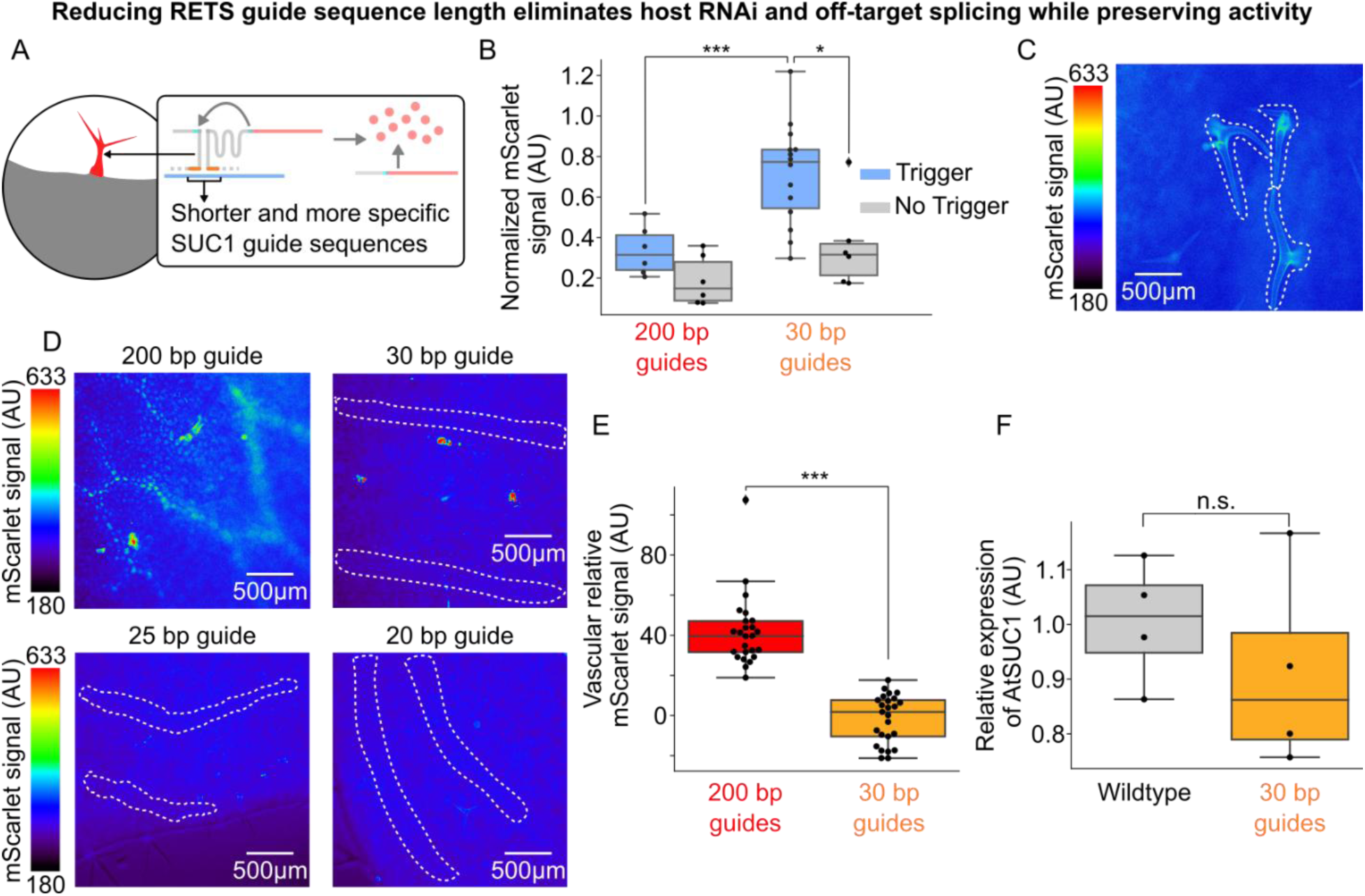
Reducing RETS guide length eliminates host RNAi and increases specificity. A) Cartoon depicting how RETS guide sequences were shortened to reduce host RNAi while preserving tissue-specific expression. B) Box plots summarizing normalized mScarlet signal from transient assays comparing SUC1:RETS:mScarlet constructs with either the original 200 bp guides (red) or new 30 bp guides directed at a new, unique, region of SUC1 (orange) either with (blue) or without (gray) the SUC1 trigger. Each dot is an independent biological replicate and * or *** correspond to p ≤ 0.05 or p ≤ 0.0005 respectively. C) Representative fluorescence microscopy images of mScarlet signal in adult leaves of a line expressing SUC1:RETS:mScarlet with shortened guides. Trichomes are highlighted with dashed white lines. D) Representative fluorescence microscopy images of mScarlet signal in the vasculature of adult leaves of plants expressing SUC1:RETS:mScarlet with either 200 bp, 30bp, 25bp, or 20bp long guide sequences. Vasculature is highlighted with dashed white lines. E) Boxplots summarizing the quantification of mScarlet signal from images of the vasculature in adult leaves of plants expressing SUC1:RETS:mScarlet with either 200 bp (red) or 30bp (orange) long guide sequences. Every dot is an independent measurement and *** corresponds to p ≤ 0.0005. F) Box plots summarizing expression AtSUC1 relative to a housekeeping gene in wildtype (gray) and SUC1:RETS:mScarlet with 30bp guide expressing plants (orange). Each dot represents a biological replicate and n.s. stands for non-significant difference.

Having determined that the shortened and more specific guide sequences were functional, we used these T-DNAs to generate stable *Arabidopsis* lines to test if these modifications reduced the RNAi and off target effects associated with the 200 bp guides. We first tested the impact of shortened guides on silencing by measuring the expression of *SUC1* in rosette leaves of the T1 progeny of these lines along with wildtype *A. thaliana* grown in parallel as described above. We observed that all the lines with the shorter guides directed to the new target sequence had no significant difference in SUC1 expression compared to the wildtype control, demonstrating that reducing the guide length eliminated the RNAi response (Figure 4F, S4 for other guide lengths).

We further characterized the expression of the mScarlet reporter in the leaves of these lines to determine whether these modified guides were able to drive trichome expression without the off-target vascular expression seen in the 200 bp long guides. We observed expression of the reporter in trichomes across the lines with shortened guides which demonstrates that this modification did not impair tissue-specific expression (Figure 4C, S5). We also observed that all the lines with RETS constructs that targeted the more specific region of *SUC1* had no visible signal in the vasculature compared to those targeting the non-specific region that did, irrespective of guide length (Figure 4D). We further quantified the difference in reporter signal from the vasculature in lines with either the original non-specific 200 bp guide sequences or the specific 30 bp guides and observed that the more specific guides reduced the off-target signal to background levels (Figure 4E). Taken together these results show that shortening RETS guide sequences can reduce RNAi as well as enhance transcript accumulation in response to the trigger, and that targeting unique regions of genes is necessary to ensure on-target tissue-specific expression.

### RETS enables studying expression patterns of genes *in planta*

Our characterization of *SUC1*:RETS:mScarlet plant lines demonstrates how this tool could be used to study the expression patterns of native genes *in planta*. To further demonstrate its utility and the ease of target programmability we designed two new RETS constructs which drive mScarlet expression in the presence of different genes (Figure 5A and 5C). We chose *PRODUCTION OF ANTHOCYANIN PIGMENT 1* (*PAP1*), a MYB transcription factor which has an intron encoded enhancer that drives its sucrose inducibility and expression in floral sepals (Broeckling et al., 2016), and *STERILE APETALA* (*SAP*), a transcription factor involved in floral identity^18^ that previous antisense hybridization studies (Byzova et al., 1999) determined is expressed in the flanks of young flower buds. Like the *SUC1* RETS constructs, the split reporter was an Omega-enhanced mScarlet, but the splice site was moved to the middle of the gene at the second U base of codon F66. This choice was made to facilitate more efficient qPCR verification of splicing and to demonstrate how the splice site could be reconfigured for future cargoes (Supplemental RETS construct design). In these constructs the guides were 50 bp long as we aimed to limit the RNAi observed with 200 bp guides but had not validated that 20 bp guides were functional yet. T-DNAs encoding expression cassettes for these split reporters as well as a constitutively expressed Venus were constructed and used to generate stable *A. thaliana* transgenic lines. The expression of the mScarlet reporter in these lines across target and non-target tissues was subsequently characterized using fluorescence microscopy.

**Figure 5.**
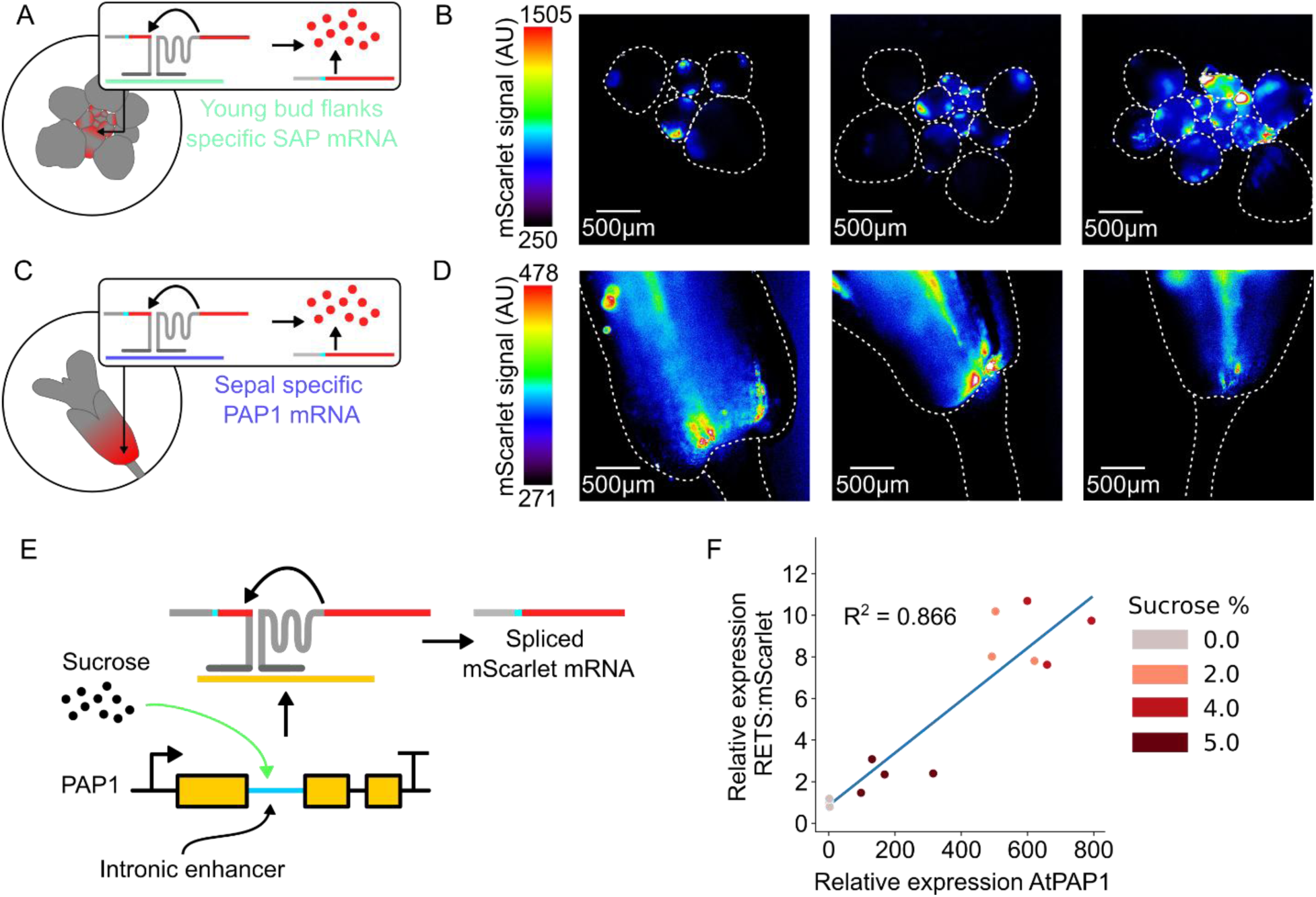
RETS can be re-targeted to other genes for dose-dependent tissue-specific expression. A) Cartoon depicting the expected expression of the mScarlet reporter in the flanks of young buds in SAP:RETS:mScarlet expressing lines, based on the previously validated expression of SAP mRNA (teal) in this tissue. B) Representative fluorescence microscopy images mScarlet signal in dissected buds of independent SAP:RETS:mScarlet lines. Dashed white lines highlight where individual buds are in the image. C) Cartoon depicting the expected expression of the mScarlet reporter in the base of sepals in PAP1:RETS:mScarlet expressing lines, based on the previously validated expression of PAP1 mRNA (purple) in this tissue. D) Representative fluorescence microscopy images mScarlet signal in dissected flower bases of independent PAP1:RETS:mScarlet lines. Dashed white lines highlight where the flower base and pedicle are in the image. E) Cartoon depicting how the intron encoded enhancer in PAP1 triggers sucrose induced expression of PAP1, thereby altering the concentration of spliced mScarlet mRNA. F) Scatter plot depicting paired measurements of spliced mScarlet mRNA and PAP1 expression relative to a housekeeping gene in PAP1:RETS:mScarlet seedlings treated with 0%, 2%, 4%, or 5% sucrose for 24hrs. Each dot is an independent biological replicate, and darker red colors represent higher sucrose concentrations.

We dissected and imaged the apexes of developing inflorescences and observed mScarlet signal only in the flanks of young developing flower buds (Figure 5B) in the *SAP*:RETS:mScarlet lines, and not in the adjacent tissues where *SAP* is not known to be expressed, demonstrating the functionality of this reporter. In the *PAP1*:RETS:mScarlet lines we imaged adult flowers and observed mScarlet signal localized in the sepals as expected and absent from adjacent floral pedicel tissues (Figure 5D). Critically, across both these constructs the two halves of the RETS system and the Venus normalization control were all driven by constitutive promoters (UBQ1 and p35s respectively) and while Venus fluorescence was observed in all cells of the plant the RETS-gated mScarlet signal was only observed in cells where the target transcript is expressed (Figure S6). This, coupled with the distinct distribution of mScarlet signal across the *SUC1*:RETS:mScarlet, *SAP*:RETS:mScarlet, and *PAP1*:RETS:mScarlet lines demonstrates that RETS can drive tissue-specific expression by piggybacking off the expression profile of a gene, independent of whether it has regulatory elements outside the canonical promoter.

The utility of RETS-based reporters as a tool to study expression patterns necessitates that the reporter signal match native gene expression in both localization and strength. To test this, we measured whether the RETS-based reconstitution of the mScarlet reporter had a dose-dependent relationship to the abundance of the target transcript by leveraging the sucrose inducibility of *PAP1* (Broeckling et al., 2016). We treated 4-day old *PAP1*:RETS:mScarlet seedlings for 24hrs with either 0%, 2%, 4%, or 5% sucrose and subsequently harvested RNA from them to enable quantification of both *PAP1* expression as well as splicing based reconstitution of mScarlet using RT-qPCR. We detected significantly higher PAP1 expression as well as correctly spliced mScarlet transcript in the sucrose treated samples as compared to the mock treated control, and the levels of RETS-spliced mScarlet correlated well with the PAP1 expression (R^2^ = 0.866) (Figure 5F). This validates the target dose dependency of the RETS-based reporters. Taken together, these results demonstrate the utility of RETS-based reporters as genetically encoded biosensors to study the expression patterns of native genes.

### RETS enables precision engineering of plant morphology

Beyond studying native gene expression patterns, the RETS system’s capacity to enable tissue-specific expression of transgenes without needing tissue-specific promoters also opens frontiers for precision engineering of plant phenotypes. As a proof-of-concept of this approach we designed a RETS construct that targets a gene with a tissue-specific expression profile and reconstitutes a developmental regulator that generates predictable changes to cell expansion and thereby organ size. We chose to target the gene, *KNOTTED*-like from *Arabidopsis thaliana* 2 (KNAT2), which is known to be expressed specifically in the floral meristem and root elongation zone based on tissue-specific RNA-seq (Fucile et al., 2011; Waese et al., 2017) (Figure 6A) so we could restrict the phenotypic alterations to these tissues. For the reconstituted developmental regulator, we chose a stabilized mutant from a family of transcriptional regulators called DELLA proteins (Davière & Achard, 2016). Prior studies have demonstrated that this mutated DELLA, called GAIrht, when expressed constitutively or from its native promoter, restricts cell expansion and hence leads to a reduction in organ size across all vegetative tissues in the plant (Willige et al., 2007). It has also been shown to inhibit the transition from vegetative to floral growth and thereby delay flowering (Li et al., 2016; Willige et al., 2007; Zhang et al., 2023). Thus, we hypothesize that this *KNAT2*:RETS:GAIrht construct would result in plants that have a reduction in root length, as the expression of GAIrht in the root elongation zone would prevent cell expansion, but no change in the size of other vegetative tissues (Figure 6A). We would also predict that plants expressing this construct would have delayed inflorescence bolting due to expression of GAIrht in floral meristems (Figure 6A).

**Figure 6.**
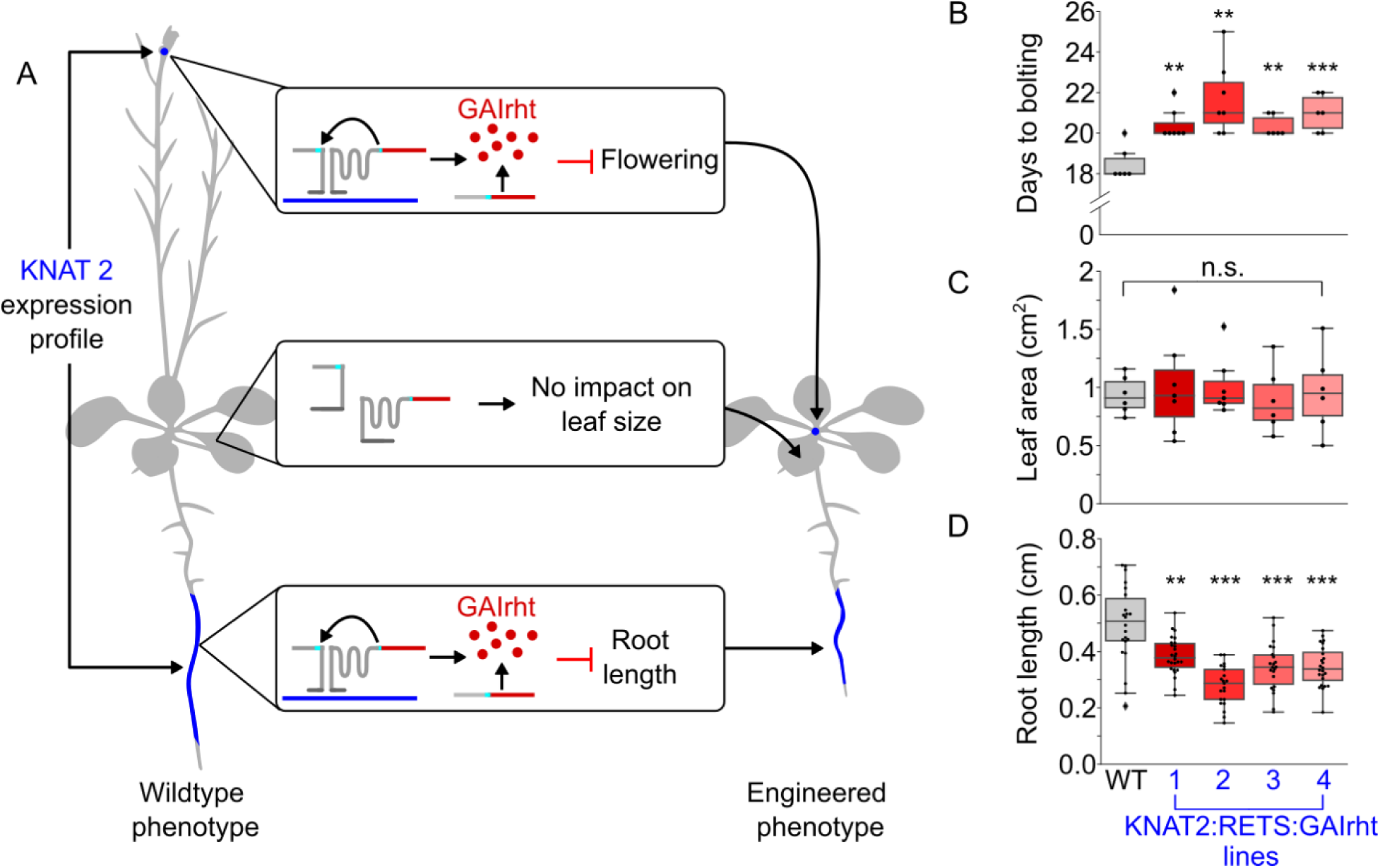
RETS enables tissue-specific engineering of plant morphology and flowering. A) A cartoon depicting that KNAT2 is expressed in the elongation zone and shoot apical meristem (blue). The middle boxes and plant image on the right show the KNAT2 dependent reconstitution of GAIrht should inhibit root cell expansion and the transition to flowering but not reduce leaf area due to the lack of KNAT2 expression in leaves. B) Box plots summarizing the flowering time, quantified as days to bolting post light shock, in either wildtype (gray) or in four independent KNAT2:RETS:GAIrht lines (reds). C) Box plots summarizing area of the same age leaf in either wildtype (gray) or in four independent KNAT2:RETS:GAIrht lines (reds). D) Box plots summarizing primary root length of seedlings either wildtype (gray) or in four independent KNAT2:RETS:GAIrht lines (reds). Each dot is an independent biological replicate and *, **, and *** correspond to p ≤ 0.05, p ≤ 0.005, and p ≤ 0.0005 respective.

A T-DNA encoding expression cassettes for each of the two halves of the *KNAT2*:RETS:GAIrht construct as well as a constitutively expressed Venus marker were constructed and used to generate transgenic *A. thaliana* lines. These lines were phenotyped in tandem with wildtype *A. thaliana* controls to assess if we observed tissue-specific changes in organ size. We observed that 3 days post germination on ½ MS agar plates *KNAT2*:RETS:GAIrht seedlings across all lines showed significantly reduced root lengths, between 0.56 and 0.78 fold, compared wildtype controls (Figure 6D). These seedlings were subsequently moved to soil and then phenotyped 7 days after the start of bolting. No significant differences were present in the *KNAT2*:RETS:GAIrht lines compared to the wildtype control in terms of developmentally equivalent leaf area (Figure 6C). This is consistent with what we would expect if RETS was successfully restricting the expression of GAIrht to tissues expressing KNAT2. We also observed that KNAT2:RETS:GAIrht plants began bolting a mean of 2.4 days later than the wildtype plants (20.9 days vs. 18.5 days) which is consistent with RETS driven expression of GAIrht in floral meristems (Figure 6B). Taken together, these results demonstrate that RETS can enable tissue-specific modulation of plant phenotypes without defined tissue-specific promoters.

## Discussion

The challenge of precise spatiotemporal control of transgene expression is often dependent on the complexity of native transcriptional regulation (Bandopadhyay et al., 2010). The canonical proximal promoter region 1-2Kb upstream of a gene’s transcription start site is often assumed to contain most of the relevant regulatory elements that drive the expression profile of a cognate gene. This assumption can provide a convenient way for synthetic biologists to choose promoter regions that express desired transgenes in specific tissues(Tian et al., 2022, 2023; Yasmeen et al., 2023), but it is becoming clear that this model is not generally applicable. Genes such as SUC1 (Lasin et al., 2020), PAP1 (Broeckling et al., 2016), and AGAMOUS (Hong et al., 2003) in *Arabidopsis thaliana* have been shown to have native expression profiles that depend on the presence of intronic enhancer elements, and more such genes are continually being discovered (Back & Walther, 2021; Meng et al., 2021; Ullah et al., 2018). Other situations such as when a crucial enhancer element is genomically distant from the affected gene (Hummel et al., 2023) or when a gene is controlled by long noncoding RNAs (Csorba et al., 2014) or microRNAs (Dong et al., 2022) that require native-sequence specificity complicate a synthetic biologist’s ability to engineer transcriptional control mechanisms that mimic native expression patterns.

Growing transcriptomic resources for model and crop plant species provide a wealth of information on the precise patterns of expression of native genes (Fucile et al., 2011; Waese et al., 2017). Our work shows that the RETS platform can leverage this information to enable spatiotemporally precise transgene expression without being confounded by cryptic regulatory motifs. The RETS platform builds on the work of previous groups that aimed to engineer the *Tetrahymena* group I intron to enable targeted and/or conditional *trans*-splicing (Alexander et al., 2005; Gambill et al., 2023; Hasegawa et al., 2004, 2006; Hasegawa & Rao, 2006; Kruger et al., 1982; Sullenger & Cech, 1994; Watanabe & Sullenger, 2000; Zaug et al., 1986). Whereas past work focused on *in vitro*, mammalian, yeast, and bacterial cell lines, our work is focused on how it can be adapted to solve a grand challenge in multicellular synthetic biology, namely spatiotemporally precise transgene expression. Our principal innovations are the elucidation of design principles for RETS to enable flexible coupling to transgenes, overcome low expression or off-target activity, and avoid host RNAi responses. We also demonstrate how these innovations make the RETS platform a useful tool for studying gene expression patterns and engineering plants.

The inclusion of the Omega leader sequence as the 5’UTR of the RETS splice product provided approximately 41% greater induction above background compared to the same construct lacking it. The likely mechanism for this increase is the Omega leader functioning to enhance the rate at which ribosomes associate with it by recruiting elongation factors (Gallie, 2002), increasing amount of protein produced. We hypothesize that the mRNA splice product created by the RETS system is relatively low abundance, so the translational enhancement is helpful to produce a robust signal. In the future the RETS system’s output strength could be improved by using stronger translational enhancers or alternative signal amplification mechanisms. A welcome byproduct of the inclusion of the omega enhancer was the creation of a generalized splice site at the U-base of the start codon ‘AUG’. Because the internal guide sequence of the ribozyme must be changed for every new splice site (Köhler et al., 1999), and each splice site has the potential to have variable efficiency (Hasegawa et al., 2006), the establishment of a consistent and functional site at the start codon allows for the modular swapping of cargoes. This contrasts with the recent work on a similar *in planta* ribozyme-based system which depended on a split GFP to provide a consistent splice site and a P2A sequence to introduce modular cargos (Y. Liu et al., 2025). In addition to reducing cargo size requirements, we hypothesize our method for enabling modular coupling would circumvent the significantly reduced expression that results from placing a cargo downstream of a P2A sequence (Z. Liu et al., 2017).

Additionally, our in-depth characterization of RETS functionality in stable transgenic lines revealed two major issues with the use of split TtRz-based systems in plants, and likely other multicellular eukaryotes, that were not apparent from the largely transient studies carried out in this recent study (Y. Liu et al., 2025). Firstly, even though longer guides have been shown to be more effective in other hosts (Ayre et al., 1999; Gambill et al., 2023; Hasegawa et al., 2006), the formation of dsRNA intermediates prior to splicing triggered an RNAi response leading to the formation of siRNAs that knockdown the RETS transcripts and/or the target sequence, drastically impacting the system’s effectiveness. This problem was evident in RETS lines with 200bp guide sequences where the expression of the target, At*SUC1*, was observed to be suppressed compared to wildtype plants. The second issue was that the association of the guides to a target did not require a 100% sequence match, the free energy of association of the two guides only needed to be sufficient to bring the two pieces together. This meant that imperfect matches, such as matches to genes closely related to the chosen target, could facilitate the splicing reaction. This issue was demonstrated in stable lines with the observation that RETS-spliced mScarlet signal was detected in the vasculature of rosette leaves, a tissue expected to have expression of the SUC1 relatives SUC2 and SUC9. Our results demonstrated that reducing guide sequence length below minimum duplex requirements for DICER proteins to efficiently generate siRNAs (Kakiyama et al., 2019) eliminated host RNAi responses and generally outperformed longer guides in terms of both fold change and absolute on-states. Additionally, this shortening enabled selection of a highly specific target region which eliminated off-target activity associated with cross targeting. However, there remain many aspects of guide sequence design that remain to be elucidated in future work, such as the precise lower limit of length that preserves functionality, as well as the impact of guide secondary structure and target binding energy on splicing efficiency.

A powerful application of RETS is the construction genetically encoded biosensors that enable studying spatiotemporal patterns of gene expression, without disrupting endogenous loci, by post-transcriptionally coupling reporter expression to transcript abundance. The tissue-specific and dose dependent expression of reporter signal in the *SUC1*:RETS:mScarlet, *PAP1*:RETS:mScarlet, and *SAP*:RETS:mScarlet lines demonstrate that RETS-reporters can be used to determine both the location and strength of gene expression. This system has several advantages over existing strategies to study gene expression. They enable non-destructive, cellular-resolution visualization of transcript abundance over developmental time in contrast to RNA-seq, RT-qPCR, or FISH, all of which necessitate sacrificing the tissue being studied, making them well suited for studying transcriptional dynamics. They can also be easily multiplexed, thanks to the ease of swapping targets and coupled transgenes, which could enable coupling different targets to spectrally separable reporters in future studies. However, these tools also have limitations, such as relatively low signal and the need to express two transcripts per target. Overcoming these challenges will be necessary to improve the utility of RETS as a tool for studying gene expression.

Beyond studying expression patterns, the tissue-specific transgene expression enabled by RETS opens the door to sophisticated plant engineering. Localizing the impact of transgene expression to specific tissues has the potential to circumvent the negative pleiotropies sometimes associated with constitutive overexpression. Our results showing that RETS enables targeted dwarfing of roots and delayed flowering without associated dwarfing in leaves, through coupling expression of a stabilized DELLA mutant (GAIrht) to the cell-types where KNAT2 is expressed, serves as a proof-of-concept of this approach. These stabilized DELLA mutants were responsible for generating cereal morphologies with semi-dwarfed shoots that drove the green revolution but also resulted in significant negative pleiotropies such as reduced nitrogen uptake by roots (Wu et al., 2021). This highlights how, while our work focuses on a model plant *A. thaliana*, the underlying technology has major translational relevance for agriculture. The broad conservation of the stabilized DELLA mutant’s impact on plant physiology (Willige et al., 2007), coupled with the host independence of TtRzs (Ayre et al., 1999; Gambill et al., 2023; Hasegawa et al., 2006; Kalvapalle et al., 2025; Y. Liu et al., 2025; Sullenger & Cech, 1994; Watanabe & Sullenger, 2000) and its easy programmability, means this strategy could be translated to crop plants in the future to facilitate trade-off free enhancement of morphology. A key limitation of this approach is that the expression of the transgene is inherently limited by the expression of the tissue-specific target. In the future, the utility of RETS for phenotype engineering could be enhanced by coupling it with secondary mechanisms that can generate a desired level of transgene expression in response to a tissue-specific trigger.

Taken together, this work demonstrates that RETS can facilitate the expression of diverse protein cargos in specific plant tissues without the need for a defined tissue-specific promoter. The key benefit of this method being the straightforward way it allows researchers to take easily accessible tissue- and single-cell level transcriptomic data and translate it into a tissue-specific expression pattern for their chosen transgene. These capabilities will make RETS will be valuable for plant synthetic biology projects that require precision timing or targeting of expression. In the future these tools could also be expanded to respond externally derived transcripts, such as viral genomes or trans-kingdom RNAs, to enable the detection and management of plant pathogens.

## Materials and Methods

### General RETS and cloning Methods

RETS constructs were designed based on vectors described in Gambill et al. (2023) (Gambill et al., 2023). The 419 base *Tetrahymena thermophila* rRNA intron ribozyme (TtRz) was split at the 15^th^ residue to produce two fragments: (1) An upstream fragment wherein the first 6 bases are the final 6 bases of the upstream exon (also referred to as the internal guide sequence binding site, IGSBS) with the 6^th^ base being a required U that defines the exon splice junction. The following 9 bases are the P1 loop of the TtRz. (2) A downstream fragment that consists of the next 11 bases of the P1 loop, a 6-base internal guide sequence (IGS) that binds complementarily to the IGSBS perfectly aside from a G that creates a wobble pairing with the splice-site U, the remaining 387 bases of the TtRz, then the downstream exon. To the 3’ end of fragment (1) a first guide sequence is added that is complementary to a target RNA of choice, preceded by helper stem sequence (GGATCA). To the 5’ end of fragment (2) a second guide sequence is added, also complementary to the target RNA, and this guide is followed by the second strand of the helper stem (TGATCC). The first and second guide sequences are chosen such that their binding sites are separated by 5 bases of target transcript to accommodate the helper stem and assist with the necessary association of the two RETS transcripts when complexed with the target RNA. Guide RNA length varied from 200 and 20 bases between experiments as noted.

Most RETS constructs in this work were designed to output the fluorescent protein, mScarlet except for the final experiments that output an autoactive rht variant of the Arabidopsis DELLA protein, GAI. Exon splice site arrangements were either designed to split the AUG start codon (utilizing the start codon’s ‘U’ base as the splice site) or a codon in the middle of the output protein’s coding sequence (UU/U of F66 in mScarlet, or CU/C of L56 in GAIrht). For each splice site, the IGS needed to be modified to match the bases in the IGSBS of the upstream exon (accounting for the required G/U wobble pairing). In the cases where the start codon served as the splice site, the final 4 bases of the omega enhancer (UUAC) completed the IGSBS. After the first experiment, all output transcripts included an omega enhancer prior to their coding sequences, whether the splice site was at the start codon or not. The two cassettes expressing the first and second RETS transcripts were driven by the Arabidopsis UBQ1 (pUBQ1) and CaMV 35s (p35s) promoters respectively, and both were terminated with the Arabidopsis HSP18.2 terminator (tHSP). In every case, a third cassette with a nuclear localized yellow fluorescent protein, venus (driven by pUBQ1, terminated by tHSP), was included for ratiometric normalization and to simplify transformant selection. These three cassettes were cloned into binary vectors using a 2-level modular golden gate cloning system (Čermák et al., 2017) to create the final plasmids for transformation into agrobacterium and subsequent use in plants.

### Agrobacterium transient Infiltration

Constructs were transformed into agrobacterium strain GV3101 using the freeze-thaw method and positive transformants were selected on LB plates with 50ug/ml kanamycin. Single isolated colonies were picked after 3 days growing at 28C. Selected colonies were used to inoculate liquid LB cultures that were grown shaking at 28C overnight (12-18 hours). Cultures were centrifuged at 2500g for 5 minutes to pellet bacteria, and the pellets were washed once with infiltration buffer (10 mM MgCl2, 10 mM 2-(N-morpholino) ethanesufonic acid (MES) pH 5.6), then resuspended in the same buffer supplemented with 100uM of acetosyringone. These bacterial suspensions were adjusted to OD600 of 1.0 and combined 1:1 with each other to facilitate codelivery of RETS_Omega+Cas9, RETS_NO-OMEGA+Cas9 and the RETS suspensions were also diluted 1:1 with acetosyringone-supplemented infiltration buffer to provide Cas9 (−) conditions with equivalent effective optical densities as the Cas9 (+) treatments (0.5 effective per construct). Diluted cultures were incubated for 2-4 hours at room temperature and then gently infiltrated into fully expanded leaves of 4-6 week old *Nicotiana benthamiana* plants using needleless syringes. After 72-96 hours, 7mm leaf disks were taken with a core borer from the infiltrated zones of each leaf and their fluorescence readings were analyzed by a Tecan Spark plate reader. Background autofluorescence was determined by analyzing buffer and Cas9-only infiltrated zones, and the mScarlet and Venus readings from corresponding RETS-infiltrated zones were assessed based on a fold change above the average background reading. Final normalized readings (Figure 2C) were determined based on a ratiometric comparison (mScarlet/Venus) between the mScarlet output (proportional to RETS-mediated splicing) and the Venus reading (constitutive, proportional to the number of cells in the leaf disk). All p values reported were calculated using a student’s T Test.

### Plant growth

When grown for seed or tissue for microscopy, *Arabidopsis thaliana* ecotype Columbia (Col-0) plants are grown in soil (3:1 ratio of Promix and turface) in a growth room or growth chamber at ∼22-24C at 60% humidity and 120-150 µmol/m²/s light with a 16/8hr day-night cycle. For transient assays, *Nicotiana benthamiana* plants were grown in Promix for 3-5 weeks in a growth chamber set to 25C, the same humidity and light conditions as *Arabidopsis*. Stable *Arabidopsis* lines were generated in the Columbia background (Col-0) using the agrobacterium floral dip method. Single isolated colonies of *Agrobacterium tumifaciens* bearing the desired binary plasmids were used to start 100mL LB cultures (supplemented with kanamycin and gentamicin) that were grown to confluence shaking overnight at 28C. Cultures were centrifuged for 10 minutes at 4000g to aspirate the growth medium and pellets were resuspended in 200mL of transformation buffer (ddH_2_0 with 5% sucrose, 0.03% Silwet L-77). Densely planted Col-0 plants that have bolted with the first flowers beginning to open are then dipped into the bacterial solution and swirled for ∼10s and allowed to dry in darkness for 24hrs. The plants are then returned to light and allowed to set T0 seed. Transgenic plants are germinated under selection by inducing light shock on antibiotic-supplemented medium. T0 seed is surface sterilized with 70% ethanol, then seeds are spread onto ½ Murashige-Skoog medium plates (0.8% w/v phytoagar) with 50mg/mL kanamycin and allowed to stratify in darkness at 4C for at least 48hrs. Plates are transferred to light for 6hrs to induce germination and then covered again for 48-72hrs at 22-25C to induce etiolation. Plates are then uncovered and over the next 3-5 days, the seedlings that survive the etiolation and produce adult leaves are considered to have passed selection and move on to further analysis. Selected lines are transferred to soil, used for experiments and/or allowed to set T1 seed. T1 plants are then either used for experiments or put through another round of selection to produce a T2 generation and so on.

### Microscope imaging

All microscope images were taken using a Leica DM5500 B upright fluorescence microscope using a CTR5500 controller system, a mercury halide lamp, and the LAS X software package. Fluorescence was resolved using mScarlet (570±20nm excitation, 609±34nm emission) and GFP (470±40nm excitation, 525±50nm emission) filter cubes for mScarlet and Venus visualization respectively. Magnification was set to 10x for every image using an HCX PL Fluotar 5x objective and a 2x ocular lens prior to image acquisition. All images were 1 second exposures, in FIM light mode with the FIM and transmission light field set to the maximum settings of 100% and 6 respectively. Images were exported as raw TIFF files and prepared for publication in imageJ. Plant material for imaging was prepared as either excised leaves (SUC1), mature flowers (PAP1), or developing flower buds (SAP), that were set on a glass slide either with a drop of 0.5% Tween-20 solution and a glass coverslip (SUC1) or imaged dry and uncovered (PAP1/SAP1).

### Plant phenotyping and imageJ analysis

For Figure 4E, microscope images (mScarlet channel, all with the same capture settings and lookup table thresholds) of leaves of SUC1:RETS:mScarlet lines with either 200 base (old target) or 30 base guides (new target) were analyzed with the ROI tool in imageJ. A background ROI was selected on a region of the leaf without observable vasculature and 5 ROIs were taken along the vascular features that were visible. The mean mScarlet signal from the background measurement was subtracted from the other 5 ROIs to produce the “Vascular relative mScarlet signal” reported in Figure 4E. For synchronized growth experiments (Figure 6), T1 KNAT2:RETS:GAIrht and Col-0 seeds were arranged in rows on ½ MS plates without antibiotics, stratified and light shocked as previously described, and placed covered, vertically in the growth chamber. After 72hrs etiolating in darkness, the plates were scanned with an office copier alongside a ruler to provide scale. The plates were then examined under a fluorescent microscope to identify which plants were true transgenics – plants with no identifiable Venus fluorescence signal in the roots were excluded from the analyses. After measurement, the seedlings were allowed to grow vertically on plates for ∼2 weeks before a subset of each genotype were transferred to soil in a growth chamber. Over the following days, each individual plant was monitored for signs of bolting. Seven days after the first sign of bolting, each plant was removed and fully dissected, with each leaf and the inflorescence being arranged in developmental order and scanned. All p values reported were calculated using a student’s T Test. All code for analyzing and plotting data, as well as all raw data, can be found at https://github.com/arjunkhakhar/RETS_paper.

### Sucrose treatments

To examine sucrose dependent induction of PAP1 and correlated induction of RETS-spliced mScarlet (Figure 5F), seedlings of Col-0 and PAP1:RETS:mScarlet were germinated on solid ½ MS media and light shocked as normal. Plants were transferred to light after 72hrs in darkness and allowed to grow for 24 more hours. Seedlings were then transferred to liquid ½ MS media supplemented with either 0%, 2%, 4%, or 5% sucrose (w/v). Seedlings in liquid media were returned to the growth chamber and allowed to incubate for 24hrs. Seedlings were then pooled, 5-8 per replicate, to create 4 replicates per treatment for RNA extraction and subsequent RT-qPCR analysis. All code for analyzing and plotting data, as well as all raw data, can be found at https://github.com/arjunkhakhar/RETS_paper.

### Expression Analysis

For relative gene expression analysis, sampling was done by harvesting at least 5 whole seedlings either 10 days post germination (for SUC1 silencing analysis: Figure 3H, 4H, and S4) or 4 days post germination (for PAP1 sucrose tests: Figure 5F). RNA was extracted using the Trizol method following manufacturer’s protocols. RNA was then treated with TurboDNase according to the manufacturer’s ‘Rigorous DNase treatment’ protocol. Synthesis of cDNA was performed with LunaScript reverse transcriptase mastermix using either the ‘Supermix’ of oligo-dT and random hexamer primers (SUC1 silencing experiment), or gene specific reverse primers to recognize PAP1, SUC1, mScarlet and the housekeeping gene PP2aa3. Fold changes in gene expression were calculated using the ddCT method. All p values reported were calculated using a student’s T Test. All code for analyzing and plotting data, as well as all raw data, can be found at https://github.com/arjunkhakhar/RETS_paper.

## Abbreviations

RETS: Ribozyme Enabled Tissue Specificity
TtRz: *Tetrahymena thermophila* Ribozyme
SUC1: Sucrose-proton symporter 1
RNAi: RNA Interference
SAP: Sterile Apetala
PAP1: Production of Anthocyanin Pigment 1
KNAT2: *KNOTTED*-like from *Arabidopsis thaliana* 2

## Author information

Research was performed at Colorado State University:

Biology Building

251 W. Pitkin Street

Fort Collins, CO 80521

United States of America

## Author Contributions

MC designed the research, performed research, analyzed data, and wrote the paper; JC performed research; VV performed research; MB performed research; HC performed research; AK acquired funding, designed the research, and revised the paper.

## Supporting Information

Supplementary Materials: Additional methods, supporting data including figures S1-S6, and tables S1-S3 listing constructs, primers, and genes used in this study (DOC).

## Acknowledgments

AK’s time and the reagents costs for this work were funded by a grant from NIGMS (5R35GM155313). MC was partially funded by a grant from the Institute of Cannabis Research at Colorado State University in Pueblo (ICR-FY23-Nachappa).

## Supplementary Materials

### Supplemental RETS construct design

For figure 2, RETS constructs were designed to detect the coding sequence of an *Arabidopsis*-codon-optimized Cas9 transcript. Guide RNA sequences were 200 bases in length, and the mScarlet cargo transcript was split at the start codon. Upstream of the start codon was either the complete 67-base omega enhancer (RETS_Omega) or just the final 4 bases of the enhancer to provide a complete IGSBS (RETS_NO-Omega). A third, target construct was prepared in parallel which expressed Cas9 under control of the *Arabidopsis* UBQ10 promoter, terminated by tHSP.

For figure 3, the RETS construct was re-designed to target exon 1 of *Arabidopsis* Sucrose transporter 1, AtSUC1. For figure 4, to reduce the likelihood of inducing dsRNA-dependent RNA silencing, RETS constructs with shorter guides were produced. Targeting the same region of AtSuc1, 3 constructs were made that utilize pairs of 30nt, 25nt, and 20nt RNA guide arms instead of the 200nt guide arms of the initial construct. In addition, 3 more constructs were designed that had 30, 25, and 20nt guide arms re-targeted to a region of AtSuc1 that is more divergent from the other members of the Suc gene family (see figure S2).

To facilitate more rapid prototyping, a constitutive copy of the AtSuc1 cDNA was cloned into a binary vector under control of the CaMV 35s promoter. This construct allowed for characterization of the new Suc1:RETS constructs in *N. benthamiana* leaves with transient expression via agroinfiltration.

For the PAP1 and SAP-detecting constructs in figure 5, the backbone of the cassettes was modified slightly: The guides were all made to be 50nt in length, and the split-site of the cargo was designed to be U197 of mScarlet, the second base of F66. This change altered the IGSBS, and thus the IGS of the TtRz was modified to be 5’-GACUGC-3’ to reflect that change, allowing for successful reconstitution of the TtRz P1 helix, facilitating the splicing reaction.

For figure 6, Autoactive DELLA protein mutant GAI(*rht*) was constructed by amplifying *Arabidopsis* GAI1 (AT1G14920) from *Col-0* without the first 55 amino acids (using codon for M56 as the new start), analogous to what was described by Willige et al (2007). The RETS construct designed to detect KNAT2 was redesigned to reconstitute GAI(*rht*), splitting the transcript at U167, and having an IGSBS and IGS of 5’-GAUUCU-3’ and 5’-GGAAUC-3’ respectively. The constitutive Venus cassette was included to facilitate selection-free identification of transformants.

**Supplemental Figure S1.**
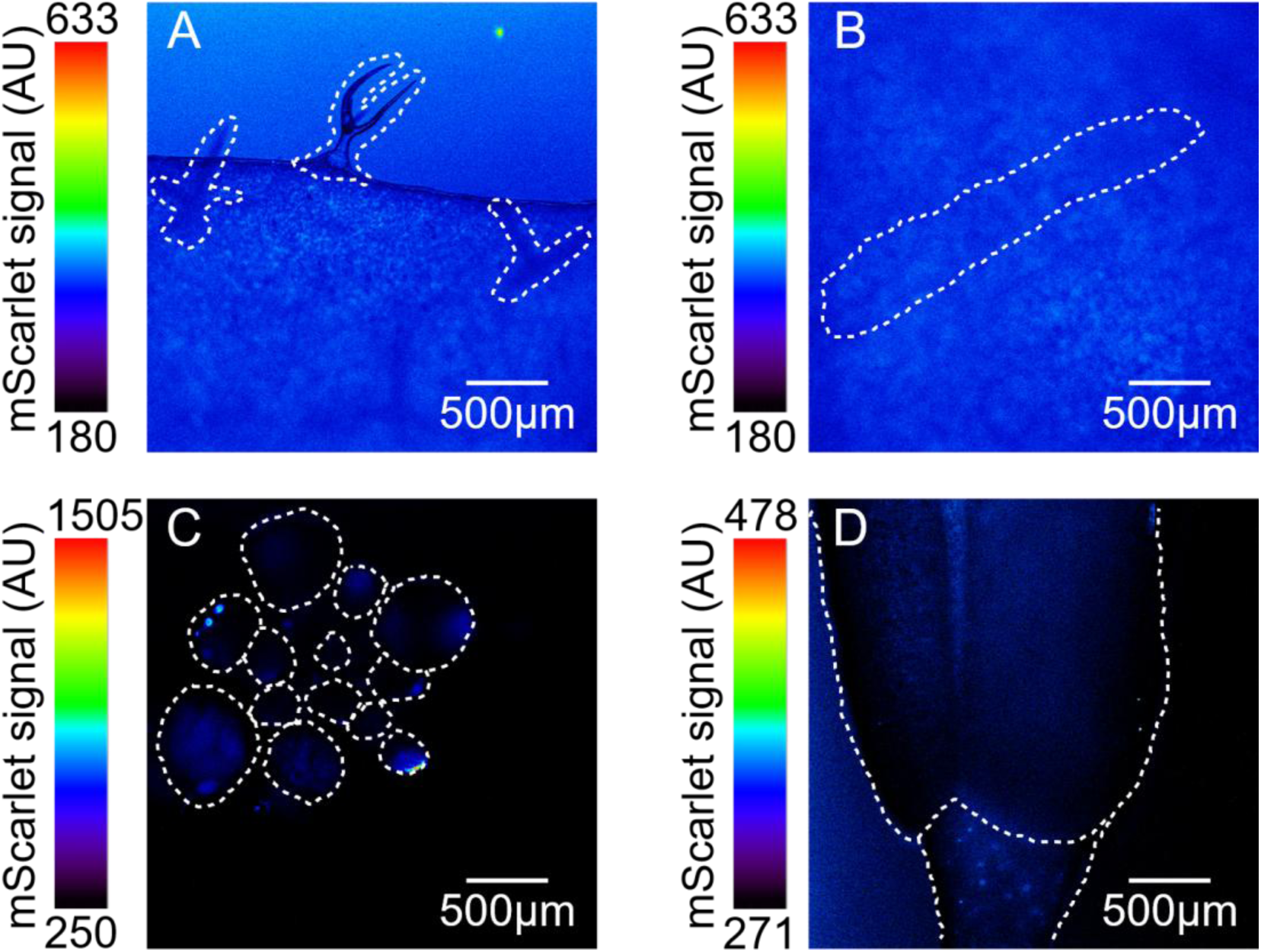
Representative *A. thaliana* Col-0 wildtype fluorescent microscope images using the mScarlet channel. A) Shows wildtype trichomes, compare to Figure 3B-D, 3F and 4C. B) wildtype vasculature, compare to Figure 4D. C) Shows developing wildtype flower buds, compare to Figure 5B. D) Shows wildtype adult flower sepals, compare to Figure 5D. Displayed lookup table thresholds are calibrated to the respective experimental images they are compared to.

**Supplemental Figure S2.**
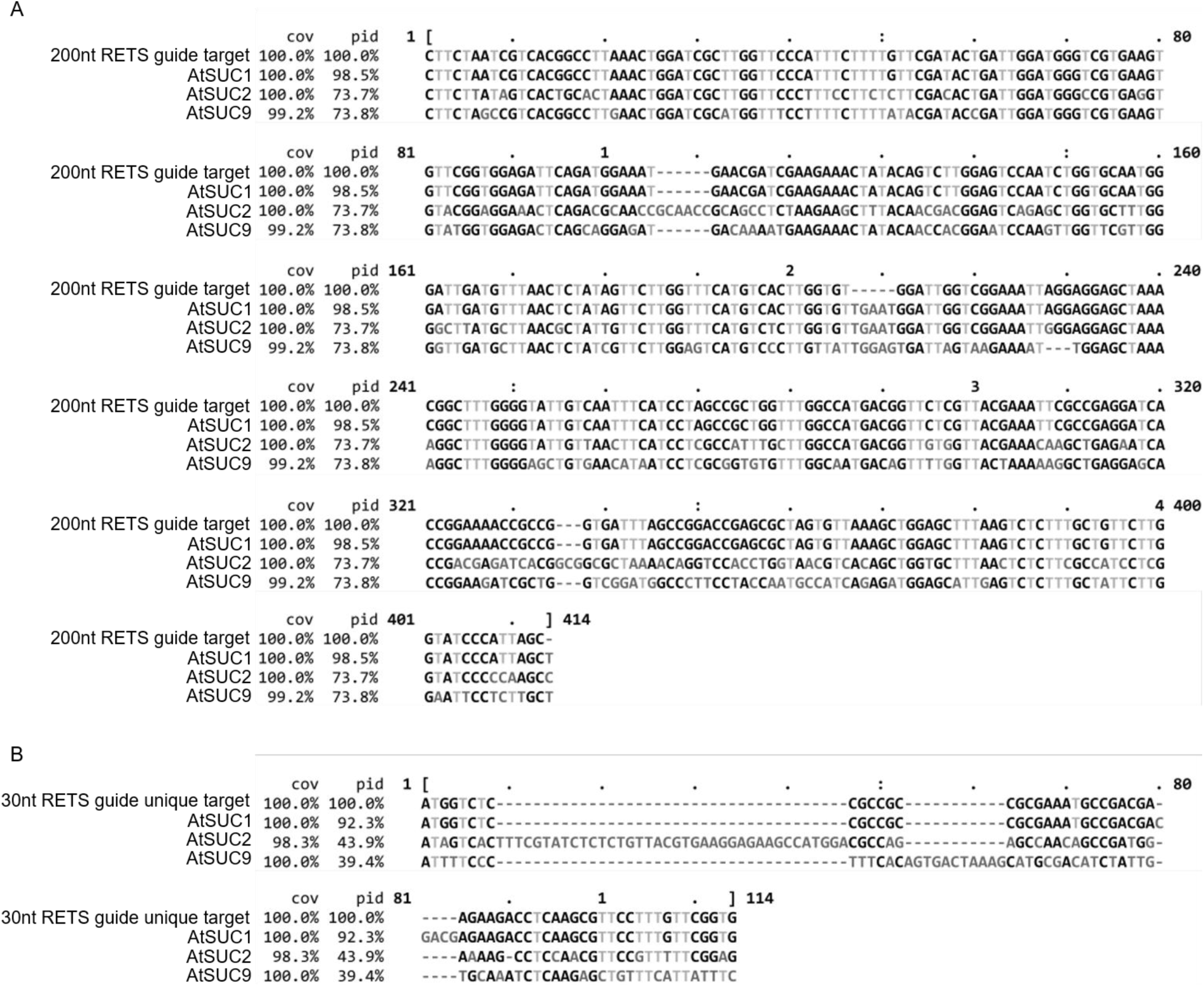
Multiple sequence alignment of RETS target sequences and SUC family genes. A) Multiple sequence alignment of the original RETS construct target sequence with 200-base guides targeting a region of AtSUC1. Target sequence is aligned to the homologous region of AtSUC1 and the major vascularly-localized members of the SUC family, SUC2 and SUC9. B) Alignment of a re-designed target site for a RETS construct with 30-base guides and SUC1, 2, and 9. Note that in both alignments, there is an intentional 5-base gap in the intended target (SUC1) not covered by the RETS guides, this allows for steric space between the two 200 or 30-base guide arms. Alignments were done using Clustal Omega and visualized with MView.

**Supplemental Figure S3.**
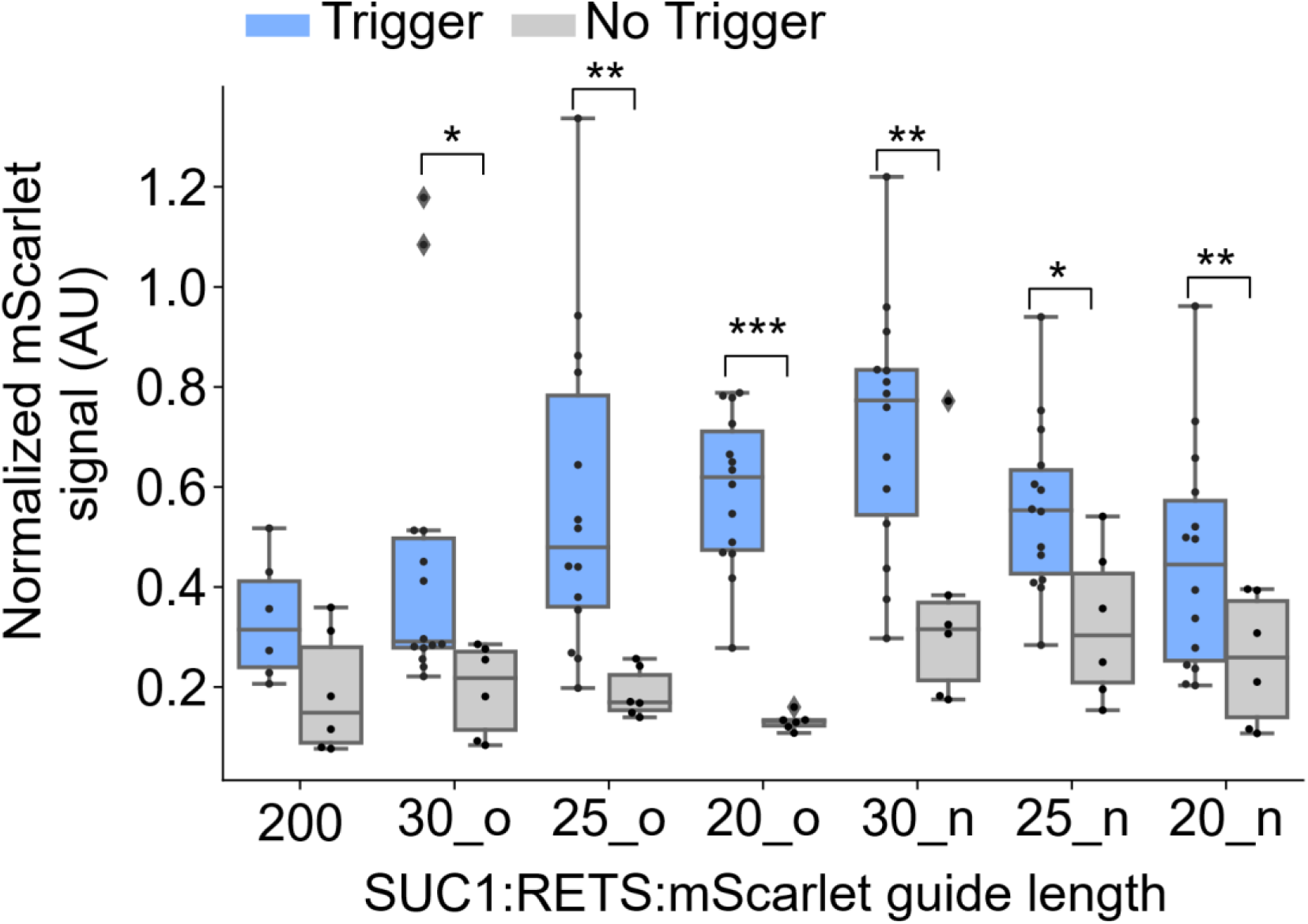
Transient expression analysis of different sized RETS guides targeting AtSUC1. Experiments were done by co-agroinfiltrating the various constructs into *Nicotiana benthamiana* leaves with or without an exogenous copy of p35S:AtSUC1 (Trigger). Blue boxplots represent data from leaf discs co-infiltrated with the trigger plasmid, grey boxplots represent leaf discs that did not have the trigger co-infiltrated with the RETS construct. Y-axis represents the normalized mScarlet fluorescence signal, i.e. the SUC1:RETS:mScarlet construct mScarlet fluorescence divided by the constitutive Venus fluorescence that was measured simultaneously. Each point represents the data from an individual leaf disc. The letter after the underscore on the x-axis label represents whether the guide sequence targets the old site (o) -the same site as the 200-base construct- or the new site (n), the site that more specifically binds to SUC1 with less cross-reactivity to SUC2/9. Statistical significance determined by Students’ t-test: * = p ≤ 0.05; ** = p ≤ 0.005; *** = p ≤ 0.0005.

**Supplemental Figure S4.**
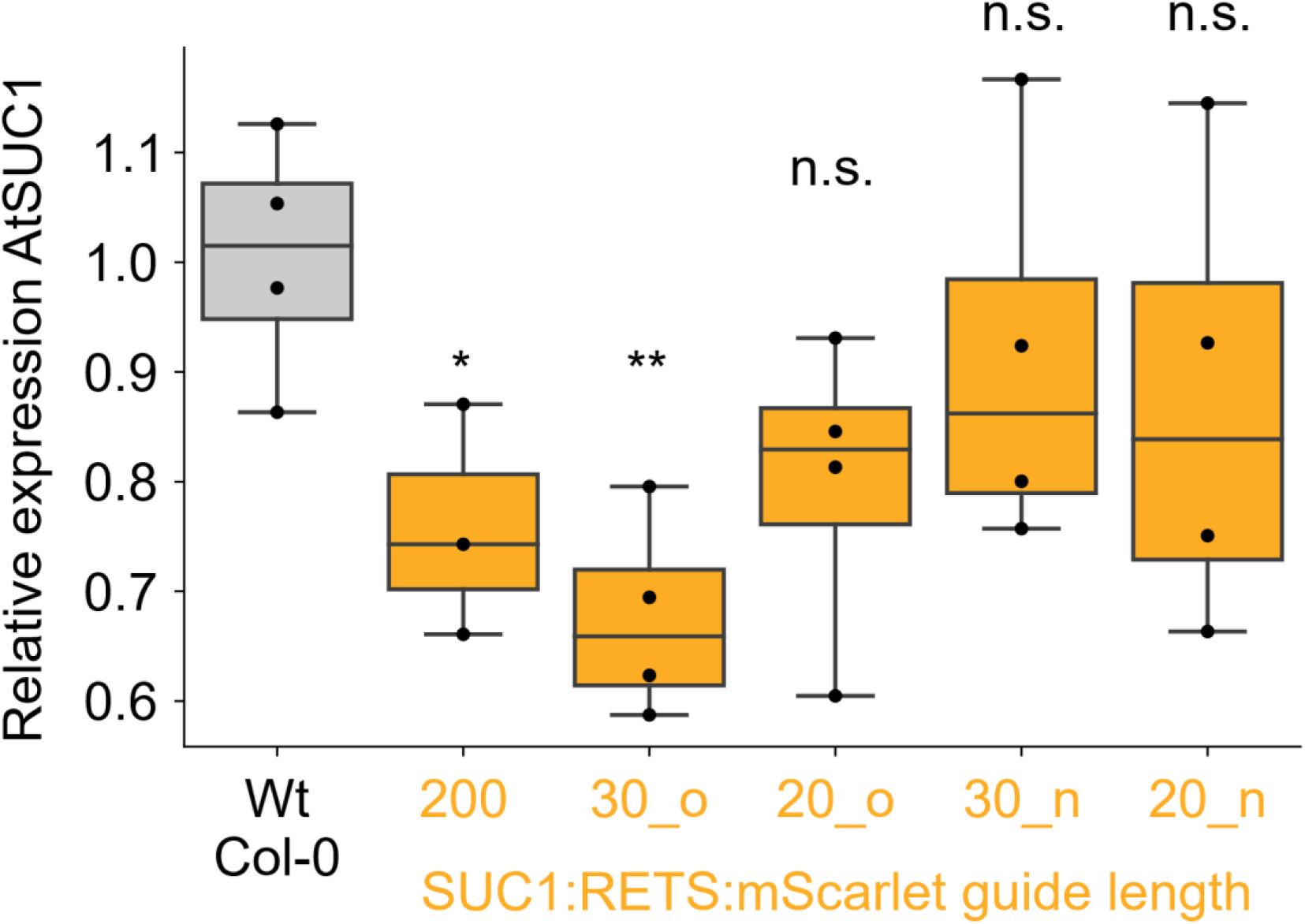
Quantitative RT-PCR measurements of AtSUC1 for seedlings of Col-0 and SUC1:RETS:mScarlet lines with different guide lengths. The grey boxplot represents the distribution of SUC1 expression in wildtype Col-0 seedlings, the orange boxplots represent the SUC1 expression from SUC1:RETS:mScarlet lines. Each point represents a biological replicate of 5-8 pooled seedlings. The letter after the underscore on the x-axis label represents whether the guide sequence targets the old site (o) -the same site as the 200-base construct- or the new site (n), the site that more specifically binds to SUC1 with less cross-reactivity to SUC2/9. Statistically significant differences to Col-0 determined by Students’ t-test: * = p ≤ 0.05; ** = p ≤ 0.005; *** = p ≤ 0.0005.

**Supplemental Figure S5.**
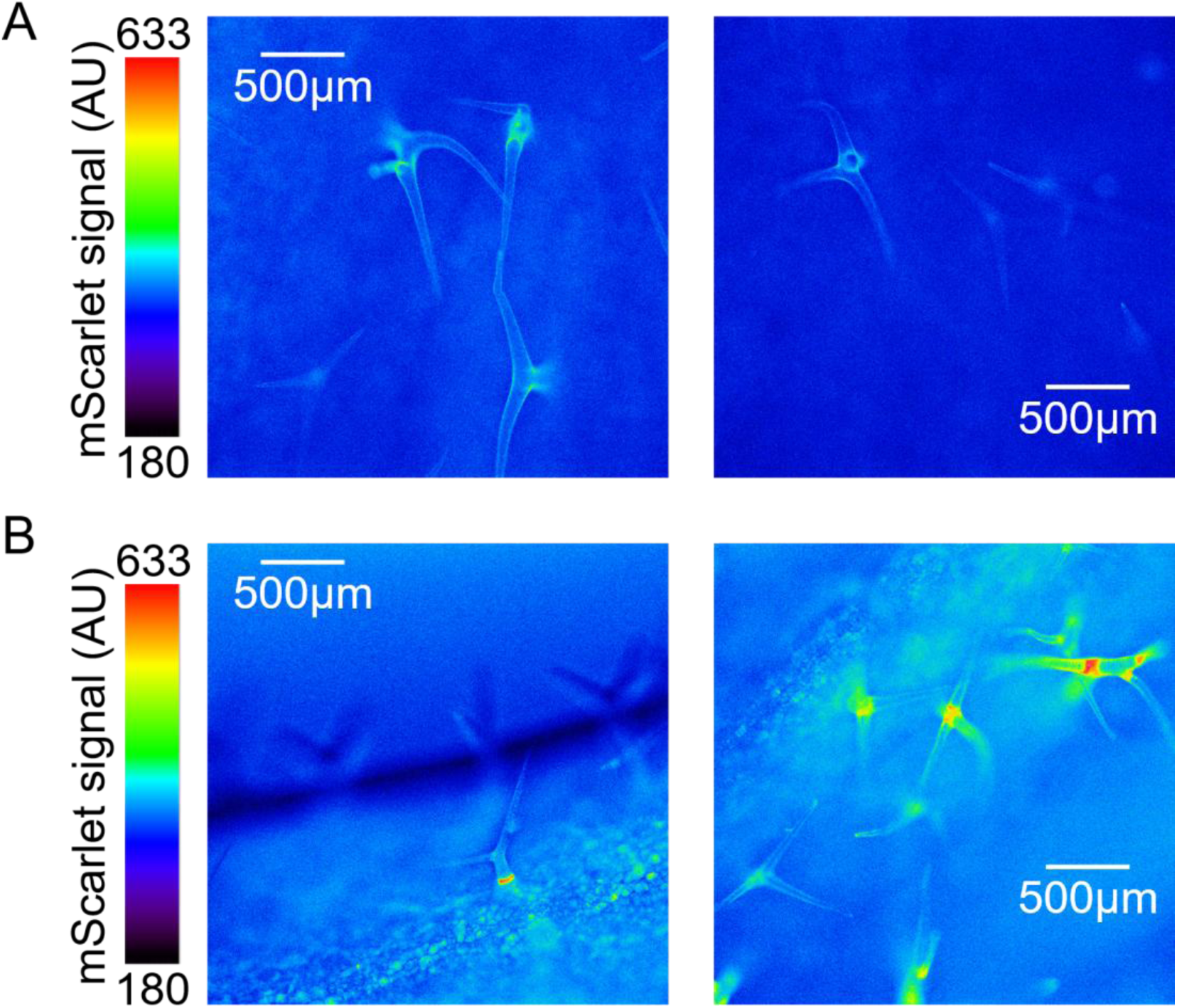
Additional microscope images of SUC1:RETS:mScarlet lines with shorter guides directed to a more unique site in AtSUC1. Images were taken of adult rosette leaves with the mScarlet channel, focusing on the trichomes. A) Images of a line with 25-base guides. B) Images of a line with 20-base guides.

**Supplemental Figure S6.**
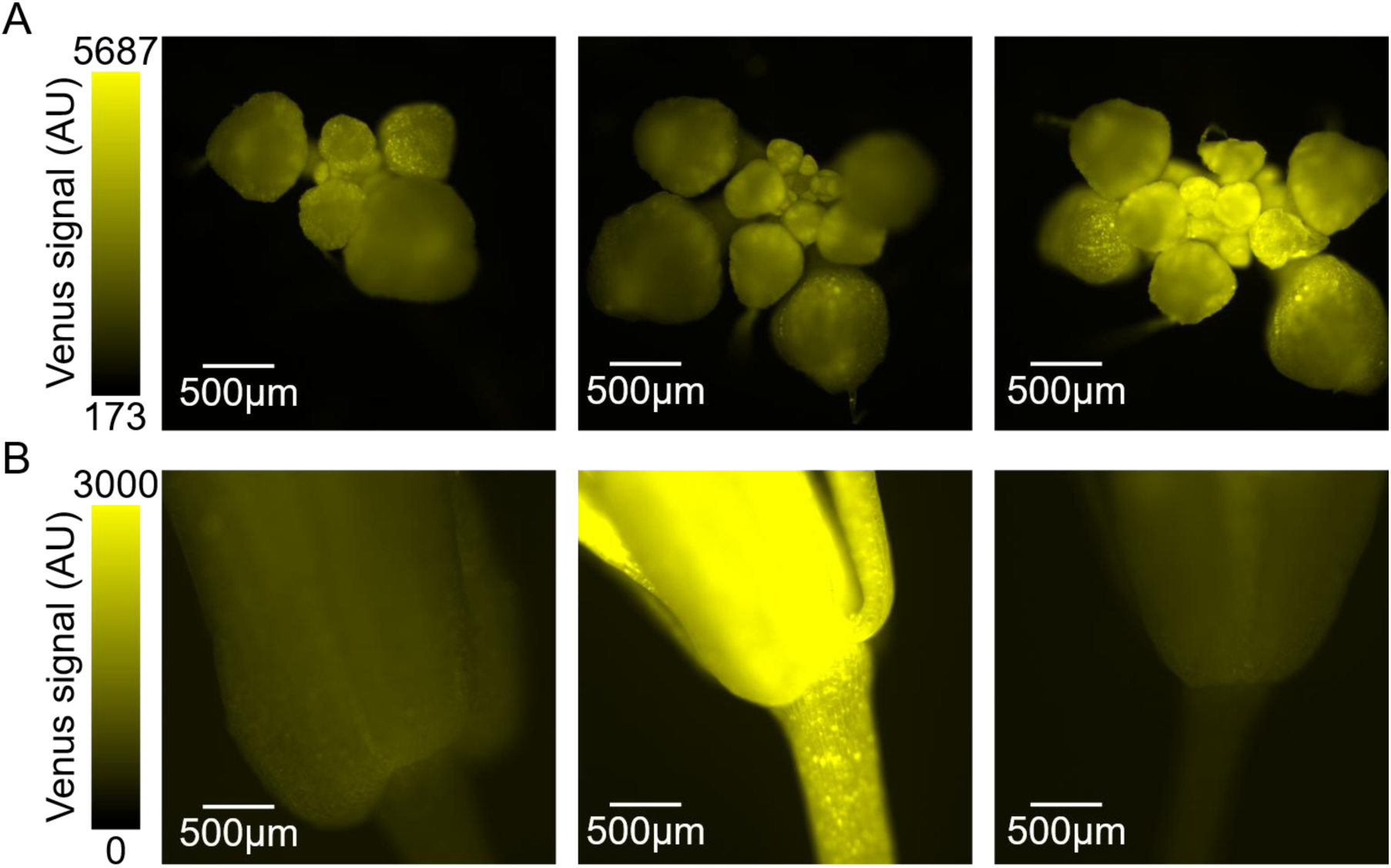
Images of SAP and PAP1:RETS:mScarlet lines in the Venus channel to show uniform distribution of the T-DNA containing the RETS cassettes. A) Developing flower buds of 3 independent SAP:RETS:mScarlet lines with constitutive Venus expression. Compare to the tissue-specific distribution of mScarlet in Figure 5B. B) Adult flower sepals of 3 independent PAP1:RETS:mScarlet lines with constitutive Venus expression. Compare to the tissue-specific distribution of mScarlet in Figure 5D.

**Supplemental Table S1.**
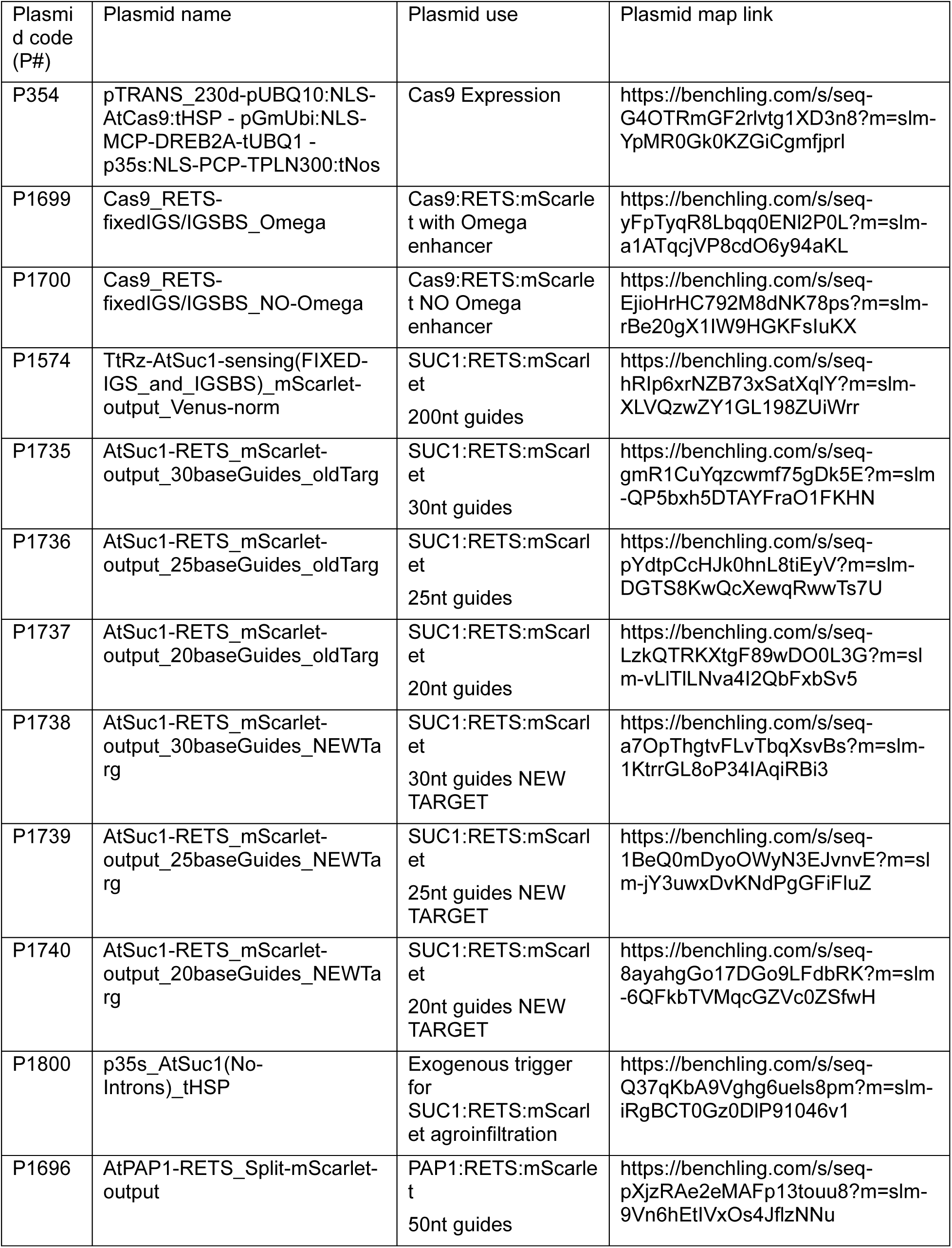
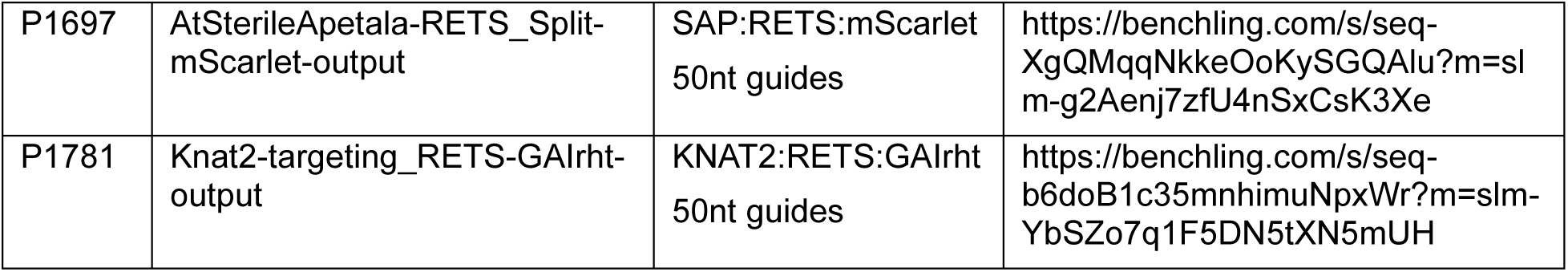
Plasmid constructs used in this work.

**Supplemental Table S2.**
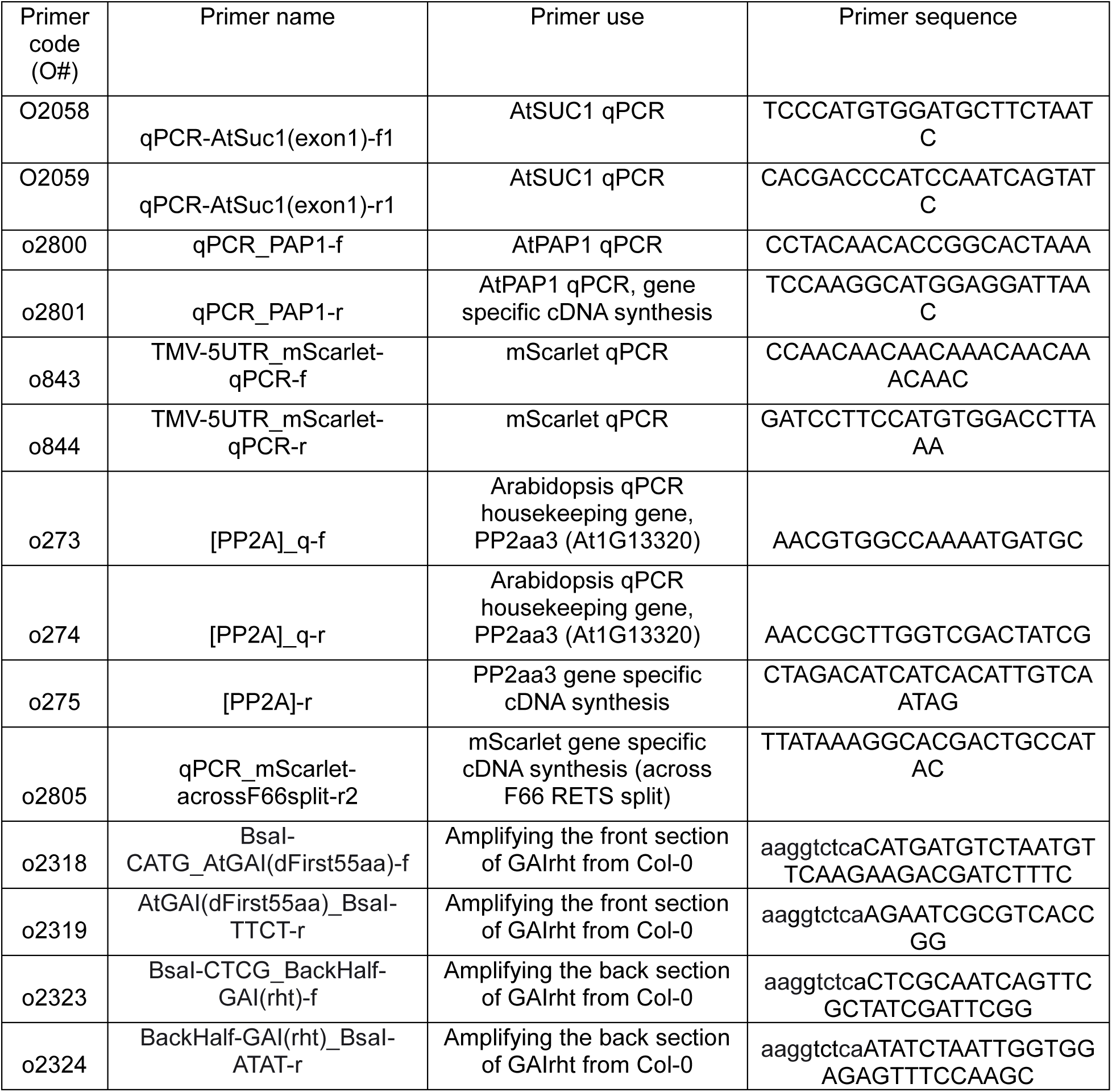
Primers used in this work.

**Supplemental Table S3.**
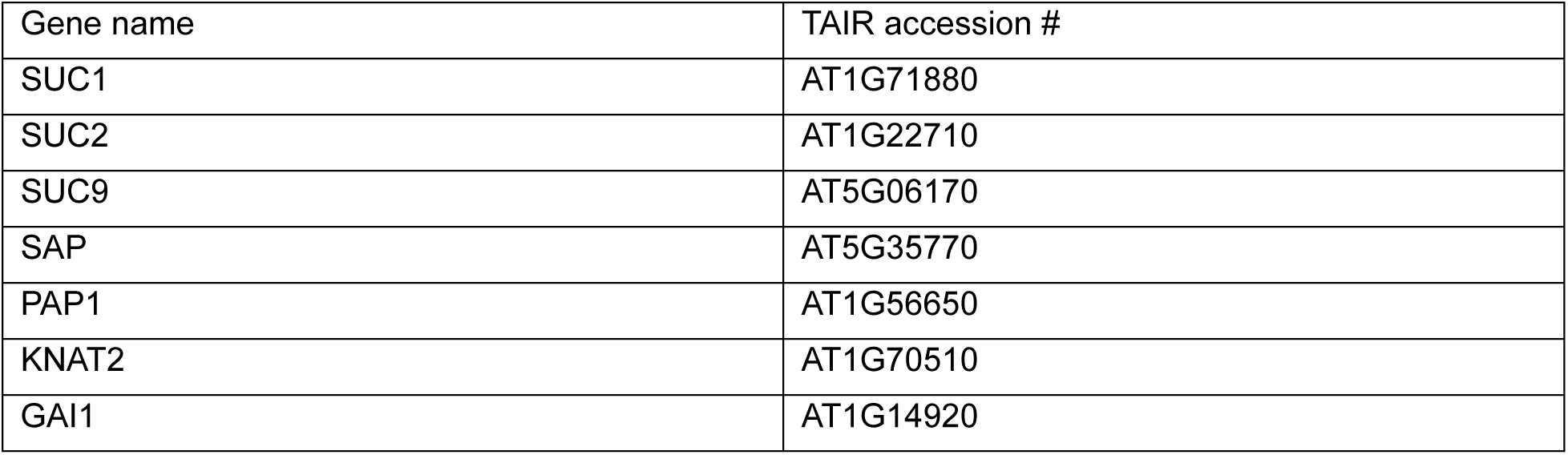
Arabidopsis target genes used in this work.

